# LiteVax-adjuvanted influenza vaccine induces transient interferon-driven innate activation and accelerates early IgG3 responses in older adults

**DOI:** 10.64898/2026.07.28.741158

**Authors:** Valentino D’Onofrio, Marjolein Verstraete, Sharon Porrez, Bart Jacobs, Azhar Alhatemi, Simon De Gussem, Sara Verwilst, Anthony Willems, Gwenn Waerlop, Fien De Boever, Ellen De Meester, Sylvie Decraene, Filip Van Niewerburgh, Geert Leroux-Roels, Elly Van Riet, Peter Paul Platenburg, Luuk Hilgers, Isabel Leroux-Roels

## Abstract

Vaccine immunogenicity declines with age due to immunosenescence, underscoring the need for improved adjuvants. LiteVax Adjuvant (LVA), a squalane-in-water emulsion containing carbohydrate fatty-acid monosulphate esters, has shown potential to enhance influenza vaccine responses, but its mechanisms in humans remain unclear. We performed a longitudinal systems vaccinology study in 56 younger (18-45 years) and older (≥60 years) adults receiving a quadrivalent influenza vaccine with or without LVA. Multi-omics analyses, including bulk and single-cell transcriptomics, flow cytometry, and serology, revealed that LVA induced a rapid, transient interferon-driven innate response, primarily in monocytes, associated with enhanced antigen processing and dendritic cell to T cell interactions. LVA promoted differentiation of influenza-specific CD4^+^ T cells toward CD45RO+ memory phenotypes, accelerated early IgG3 antibody responses, and increased IgG-producing plasmablast frequencies without affecting overall antibody magnitude. Importantly, LVA reduced age-related differences in early immune responses, improving response quality and supporting vaccine performance in older adults.

## Introduction

Seasonal influenza can cause mild or severe respiratory illness, resulting in hospitalization and, in some cases, death. Older adults, children under 5 years of age, individuals with chronic illnesses, and pregnant women are at increased risk of developing severe disease and influenza-related complications (1). Annual vaccination remains the most effective intervention and the primary strategy to prevent serious complications and death caused by influenza. However, current influenza vaccines are not affordable for many people in low- and middle-income countries, and in addition, vaccine effectiveness and immunogenicity are suboptimal in older adults (2). The effectiveness of standard-dose influenza vaccines in older adults is generally moderate and varies considerably across influenza seasons, with estimates against laboratory-confirmed influenza infection often below those observed in younger adults (3). Moreover, both humoral and cell-mediated immune responses to influenza vaccination have been shown to be reduced in older adults, including reports of up to 8-fold lower hemagglutination inhibition (HI) antibody titers (4), and reduced memory CD4+ T cell responses (5).

Aging is associated with a profound dysregulation of the immune system, resulting in impaired magnitude, quality, and durability of vaccine-induced responses (2). In addition to age, factors such as comorbidities, frailty, inflammaging, and cumulative antigenic exposure throughout the life course further influence vaccine responsiveness (6). Inflammaging, defined as a state of chronic low-grade inflammation, and immunosenescence, which reflects the complex remodeling and functional decline of the immune system, are key hallmarks of immune aging (7). Older adults experience persistent inflammatory activity while simultaneously demonstrating a reduced capacity for pathogen recognition and impaired IFN*γ* and TNFα signaling, thereby hindering efficient antigen presentation (8). Aging is also associated with an increased frequency of exhausted CD4+ T cells exhibiting reduced CD40L expression, which is required for efficient B cell activation. Similarly, B cells from older adults display a reduced capacity for immunoglobulin (Ig) class-switch recombination (7, 9).

To improve protection against influenza in older adults, several countries have incorporated recommendations for the use of high-dose or adjuvanted vaccines for older adults (10). MF59-adjuvanted influenza vaccines have shown a greater relative effectiveness against influenza-related medical encounters than standard-dose vaccines in this population (11). Adjuvants can enhance vaccine-induced immune responses through diverse mechanisms, including activation of innate immune signaling pathways, recruitment and activation of immune cells, and enhanced antigen uptake and presentation. (12). In general, adjuvants combine stimulation of pathogen recognition receptor (PRR) signaling pathways with enhanced antigen delivery. This results in robust monocyte activation, increased cytokine and chemokine production, recruitment of immune cells, including neutrophils and antigen-presenting cells (APCs), and more efficient antigen uptake and presentation (12). These processes can augment both humoral and cell-mediated immune responses and may ultimately improve vaccine effectiveness, although their mechanisms of action in humans remain incompletely understood. Insights in how adjuvants shape immune responses is crucial for the rational design of vaccines and optimization of vaccination strategies. It will inform design strategies both for antigen-sparing approaches aiming to making it more affordable for people in low- and middle-income countries, and for older adults who exhibit reduced vaccine immunogenicity.

LiteVax Adjuvant (LVA) is a novel adjuvant, that immobilizes synthetic carbohydrate fatty-acid monosulphate esters (CMS) on the surface of oil droplets of a squalane-in-water nano-emulsion. A preclinical study has demonstrated that single-dose influenza vaccines containing CMS induces antibody titers comparable to or exceeding those elicited by two doses of MF59-adjuvanted vaccines (13). Furthermore, a first-in-human study showed that 1/5 antigen dose combined with LVA induced similar antibody titers, indicating cost reduction could be feasible (14). Mechanistically, *in vitro* studies have shown that LVA activates APCs, including dendritic cells (DCs) capable of T helper cell polarization (15). Current evidence suggests that LVA may act through complimentary mechanisms, including both enhanced antigen endocytosis mediated by the oil-in-water emulsion and activation of toll-like receptors (TLR)2-and TLR4-dependent pathways by CMS (16). In a phase 1 clinical trial, addition of 1 mg LVA to a standard-dose seasonal influenza vaccine enhanced HI antibody responses, particularly against influenza A strains (17). However, the immunological mechanisms underlying these effects in (older) adults remain incompletely understood.

In this study, we use a multi-omics approach on samples from a clinical trial in which a full dose influenza vaccine was used with or without LVA, with the goal to increase immune responses in older adults (17). To study the mechanism of action of LVA in more detail, the immune responses induced by LVA-adjuvanted influenza vaccination in both younger and older adults were investigated. We show that LVA is associated with enhanced IFN-signaling in monocytes and increased antigen presentation by dendritic cells, accompanied by augmented activation of memory CD4+ T cells. Together, these responses are associated with a distinct immune profile characterized by enhanced early IgG class-switching towards the IgG1 and IgG3 subclasses and attenuation of age-associated differences in earlt transcriptional responses.

## Results

### LVA induces robust innate immune activation associated with systemic interferon responses

To define the mechanisms underlying LVA-mediated immune enhancement, we performed a longitudinal systems level analysis of immune responses following vaccination with a quadrivalent inactivated seasonal influenza vaccine administered alone (QIIV) or in combination with LVA (**Figure 1A**). A total of 56 participants were included, of whom 28 received QIIV + 1mg LVA (QIIV + LVA) and 28 received QIIV alone. Each vaccine group consisted of 12 participants classified as younger adults (18-50 years old) and 16 participants as older adults (60 years or older). Characteristics of this study population have been described previously (16). Blood samples were collected at multiple time points to capture transient inflammatory and transcriptional programs known to shape subsequent adaptive immune responses. Single-cell RNA sequencing analyses were performed exclusively in younger adults.

**Figure 1.**
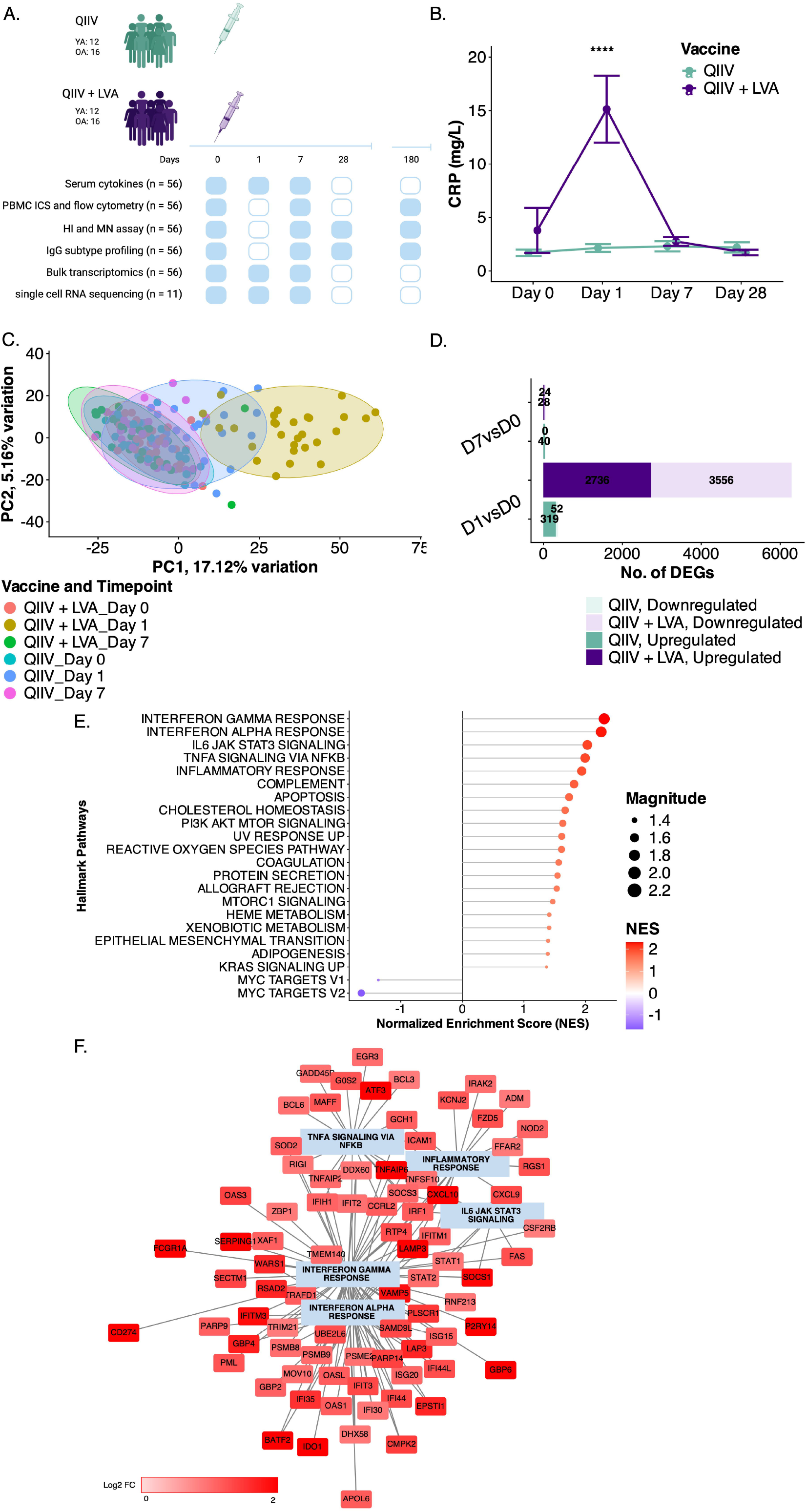
Systemic inflammatory and transcriptional responses following QIIV + LVA. (A) Study design and sampling scheme. Younger adults (YA) and older adults (OA) received quadrivalent inactivated influenza vaccine (QIIV) alone or in combination with 1mg LVA. Blood was collected at days 0, 1, 7, 28, and 180 post-vaccinations for multiomic analyses, including serum cytokines, intracellular cytokine staining (ICS) and flow cytometry, hemagglutination inhibition (HI) and microneutralization (MN) assays, IgG and IgG subclass profiling, bulk transcriptomics, and single-cell RNA sequencing in a subset of participants. (B) Mean + SEM of serum C-reactive protein (CRP) levels over time following vaccination. CRP is transiently induced post-vaccination at day 1, with a significantly higher increase observed at early time points in the QIIV + LVA group compared to QIIV alone, indicating enhanced systemic inflammation. (C) Principal component analysis (PCA) of bulk transcriptomic profiles. Separation along principal components (PC1: 17.12% variance; PC2: 5.16%) reflects clustering by vaccine formulation and timepoint, highlighting distinct early transcriptional responses induced by QIIV + LVA compared with QIIV alone. (D) Differential gene expression analysis (DEA) comparing post-vaccination timepoints (day 1 and day 7 vs. baseline (day 0)). Bar plots show the number of upregulated and downregulated genes in each vaccine group, with QIIV + LVA inducing a more pronounced transcriptional response than QIIV alone. (E) Gene set enrichment analysis (GSEA) of Hallmark pathways at day 1 post-vaccination between vaccine groups. Normalized enrichment scores (NES) reveal strong induction of innate immune pathways following vaccination, including IFN type I and type II responses, TNFα signaling via NF-κB, and IL-6/JAK/STAT3 signaling in the QIIV + LVA group. (F) Gene network representation of differentially upregulated genes after QIIV + LVA associated with key innate immune pathways. Nodes represent genes, with color indicating direction and magnitude of regulation (log_2_ fold change), and connections denote known functional or regulatory interactions. The network highlights coordinated upregulation of interferon-stimulated genes (e.g., IFIT1–3, ISG15, OAS family, RSAD2), inflammatory mediators (e.g., TNFAIP6, ICAM1, CXCL9/10), and pathway regulators linked to interferon signaling, illustrating an integrated antiviral and inflammatory response induced by vaccination. Significance levels are indicated as * (p < 0.05), ** (p < 0.01), *** (p < 0.001), **** (p < 0.0001), unless otherwise specified. CRP: C-reactive protein; HI: hemagglutination inhibition; ICS: intracellular cytokine staining; LVA: LiteVax Adjuvant; MN: microneutralization; NES: normalized enrichment score; OA: older adults (60 years or older); PBMC: peripheral blood mononuclear cells; QIIV: quadrivalent inactivated seasonal influenza vaccine; YA: younger adults (18-50 years);

We first assessed systemic inflammation by quantifying circulating C-reactive protein (CRP) concentrations. Co-administration of LVA with QIIV induced a strong but transient increase in CRP, peaking on day 1 and returning towards baseline by day 7. CRP concentrations at day 1 were significantly higher in the QIIV + LVA group than in the QIIV-alone group (15.14±16.56 mg/L versus 2.15±1.90 mg/L, respectively; p < 0.0001) (**Figure 1B**) and this pattern was observed across all participants in both age groups (**Extended Data Figure 1A**). The absence of fever, despite elevated CRP levels, suggests that LVA induces measurable systemic inflammation without a concomitant increase in body temperature (**Extended Data Figure 1B**).

To investigate the molecular programs underlying this response, we profiled whole blood (bulk) transcriptomes longitudinally. Principal component analysis (PCA) revealed a marked temporal shift in the QIIV□+□;LVA group, with the most pronounced shift from baseline observed on day 1, indicative of a rapid transcriptional response that was less pronounced following QIIV alone (**Figure 1C**; **Extended Data Figure 1C**). Consistent with this finding, differential expression analysis identified substantially more differentially expressed genes (DEGs) in the QIIV + LVA group, with the response peaking on day 1 (6292 DEGs versus 371 DEGs) and largely resolving by day 7 (52 DEGs versus 40 DEGs) (**Figure 1D**). These findings suggest that LVA induces a robust and transient innate immune response. Pathway enrichment analysis revealed that these transcriptional changes were dominated by innate immune and inflammatory signaling pathways, including TNFα signaling via NF-κB (Normalized Enrichment Score (NES): 1.99, p < 0.0001) and IL-6/JAK/STAT3 signaling (NES: 1.99, p < 0.0001) (**Figure 1E**; **Extended Data Figure 1E-F**). Notably, interferon-associated pathways emerged as the most prominently enriched pathways following QIIV + LVA administration, with significant enrichment of both type I and type II interferon responses (NES: 2.25, p < 0.0001, and NES: 2.29, p < 0.0001, respectively). This was accompanied by robust upregulation of canonical interferon-stimulated genes, including ISG15, MX1, IFIT3, STAT1, and STAT2, specifically in the QIIV□+□LVA group at day 1 (**Extended Data Figure 1D**).

To further resolve the interplay of these transcriptional responses, we constructed a gene co-expression network using genes upregulated following vaccination. This analysis revealed highly interconnected modules that were strongly enriched for interferon-stimulated genes in the QIIV□+□LVA group on day 1 (**Figure 1F**). Central nodes within this network included key regulators of interferon signaling, such as STAT1 and IRF1, both of which exhibited high connectivity, suggesting coordinated regulation of the interferon response. In contrast, genes associated with inflammatory pathways, including NF-κB and IL-6/JAK/STAT3 signaling, formed more peripheral clusters. Importantly, the IFN*γ*-associated module displayed the greatest overall connectivity and magnitude of regulation, indicating that it represents the dominant transcriptional program of the early response to LVA.

Together, these findings indicate that LVA induces a rapid but transient systemic innate immune response characterized by a central interferon signaling axis. These data further show that LVA enhances innate immune responses by establishing an early interferon-driven environment.

### Interferon responses are enriched in monocytes and associated with antigen-presentation signatures in dendritic cells

To define the cellular origin and functional consequences of the interferon-dominated transcriptional program identified by bulk RNA sequencing, we performed longitudinal single-cell RNA sequencing of peripheral blood mononuclear cells (PBMCs) collected following vaccination of younger adults. After stringent quality control and filtering (**Extended Data Figure 2**), high-quality transcriptomes were retained across all samples, with consistent sequencing depth and gene detection rates.

**Figure 2.**
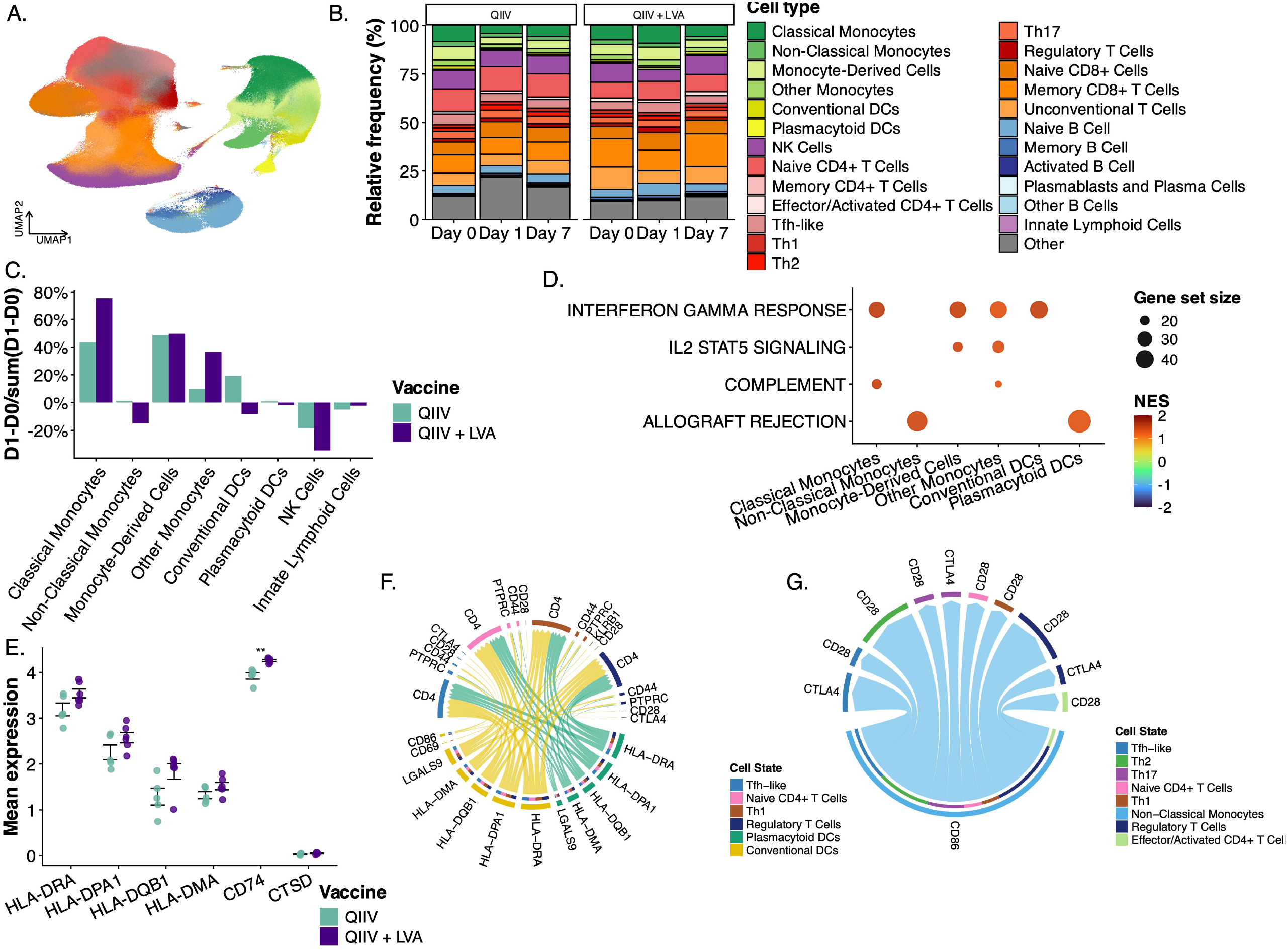
Single-cell landscape of immune responses following QIIV + LVA vaccination in younger adults. (A) UMAP embedding of single-cell RNA-sequencing data from peripheral blood mononuclear cells (PBMCs), colored by annotated immune cell types. Major populations include monocyte subsets, dendritic cells (DCs), NK cells, T cell subsets, B cell subsets, and innate lymphoid cells. (B) Relative frequencies of immune cell populations across timepoints (days 0, 1, and 7) and vaccine groups (QIIV vs. QIIV + LVA). Stacked bar plots show limited shifts in total immune cell composition following vaccination. (C) Changes in innate cell subset frequencies between day 1 and baseline (D1 minus D0, normalized to total change). Bar plots highlight preferential expansion (positive values) or contraction (negative values) of specific cell populations, with limited differences between QIIV and QIIV + LVA groups. (D) Gene set enrichment analysis (GSEA) of Hallmark pathways at day 1 post-vaccination between vaccine groups across monocytes and DCs. Normalized enrichment scores (NES) for significant pathways are shown and indicate enrichment for QIIV + LVA in red or enrichment in QIIV alone in blue. Dot size indicates the gene set size. (E) Mean + SEM of expression of antigen presentation-related genes across DC populations. No difference in the expression of MHC class II genes (HLA-DRA, HLA-DPA1, HLA-DQB1, HLA-DMA) and a significant increase of associated molecules (CD74) suggests enhanced antigen-presenting capacity following vaccination. (F) Chord diagram representing upregulated predicted cell-cell communication networks between DC populations (yellow and green) and CD4+ T cell populations after QIIV + LVA vaccination. Arcs connect interacting cell types, with edge thickness indicating the relative strength or number of inferred ligand-receptor interactions. The plot highlights consistent coordinated activation of innate and adaptive immunity. (G) Chord diagram representing downregulated predicted cell-cell communication networks between non-classical cell populations (light blue) and CD4+ T cell populations after QIIV + LVA vaccination. Arcs connect interacting cell types, with edge thickness indicating the relative strength or number of inferred ligand-receptor interactions. The plot highlights regulatory communication pathways. Significance levels are indicated as * (p < 0.05), ** (p < 0.01), *** (p < 0.001), **** (p < 0.0001), unless otherwise specified. LVA: LiteVax Adjuvant; NES: normalized enrichment score; QIIV: quadrivalent inactivated seasonal influenza vaccine;

Unsupervised clustering identified all major immune cell populations, including monocyte subsets, DCs, T cells, B cells, and innate lymphoid populations, with comparable baseline cellular composition between vaccine groups (**Figure 2A**). Cell-type proportions remained largely stable over time (**Figure 2B**). No statistically significant changes in the proportion of innate immune cell populations were observed between day 0 to day 1. However, a trend towards a greater relative increase in classical monocytes was observed following QIIV + LVA compared with QIIV alone (75% versus 44%, respectively, n.s.) (**Figure 2C**). These findings indicate that the transcriptional changes observed at the bulk level predominantly reflect functional reprogramming within immune cell populations rather than alterations in cellular composition.

We next sought to identify the cell populations contributing to the interferon signature. Pathway analysis at single-cell resolution revealed that IFN*γ* responses were significantly enriched in monocyte populations in the QIIV□ +□LVA group at day 1, including classical monocytes (NES: 1.56, p = 0.005), non-classical monocytes (NES: 1.36, p = 0.019), and monocyte-derived cells (NES: 1.49, p = 0.001) (**Figure 2D**). This enrichment reinforces the dominant interferon response detected by bulk transcriptome analyses and identifies monocytes as the principal cellular compartment associated with this signature. Other innate immune pathways, such as IL-2/STAT5 signaling and complement activation, showed more modest enrichment, further underscoring dominant interferon-driven response.

Given the central role of monocytes and DCs in antigen presentation, we next investigated whether this interferon-driven activation state was accompanied by enhanced antigen presentation capacity. LVA administration did not increase the expression of MHC class II genes (HLA-DRA, HLA-DPA1, and HLA-DQB1) in DCs (**Figure 2E**). However, expression of the antigen processing component CD74 in DCs on day 1 was significantly higher following QIIV + LVA than following QIIV alone (mean expression 4.26±0.05 versus 3.92±0.16, respectively; p = 0.026)

To determine how these transcriptional changes translate into intercellular communication, we performed ligand–receptor interaction analysis using CellChat. Chord diagram visualization revealed a marked increase in predicted signaling interactions from DCs to CD4+ T cells in the QIIV□+□LVA group compared with QIIV alone (**Figure 2F**). These interactions were enriched for pathways involved in antigen presentation and T cell co-stimulation, including MHC-II - TCR and CD86 - CD28 axes, indicating enhanced capacity of antigen-presenting cells to engage CD4+ T cells. Notably, CTLA4 also emerged as a relevant predicted receptor that interacts with dendritic cell-derived ligands, suggesting the concurrent induction of regulatory feedback mechanisms that modulate T cell activation. In contrast, non-classical monocytes contributed more prominently to the outgoing signaling network in QIIV alone compared to QIIV + LVA (**Figure 2G**).

Collectively, these findings support a model in which LVA-induced interferon signaling in monocytes is associated with molecular features consistent with enhanced antigen presentation, including increased CD74 expression and predicted strengthening of antigen-presenting cell to T cell communication networks.

### LVA promotes T helper polarization and activation of CD45RO+ CD4+ T cells

To assess whether LVA-adjuvanted vaccination was associated with changes in antigen-specific T cell responses, we characterized influenza-specific CD4+ T cell activation and differentiation using intracellular cytokine staining (ICS) and single-cell transcriptomics.

We first quantified influenza-specific CD4+ T cell responses following stimulation with peptides derived from the vaccine antigens (**Extended Data Figure 3A**). Both QIIV and QIIV + LVA induced increases in the frequencies of cytokine-producing influenza-specific CD4+ T cells across all four influenza vaccine strains as compared to pre-vaccination. Polyfunctional CD4+ T cells co-expressing CD40L and at least two cytokines increased by day 7. Although these responses gradually declined thereafter, they remained above baseline through day 180 indicating durable functional activation (**Figure 3A**). No statistically significant differences in the frequencies of polyfunctional CD4+ T cells were observed between vaccine groups at any time point. Analysis of cytokine expression patterns among influenza-specific CD40L+ CD4+ T cells demonstrated that both vaccines predominantly induced IFN*γ*- and TNFα-producing cells (**Extended Data Figure 3B**). As the CD4+ T cell responses to the four influenza strains exhibited similar kinetics, subsequent analyses were performed using the mean response across strains as a summary measure of the overall influenza-specific response.

**Figure 3.**
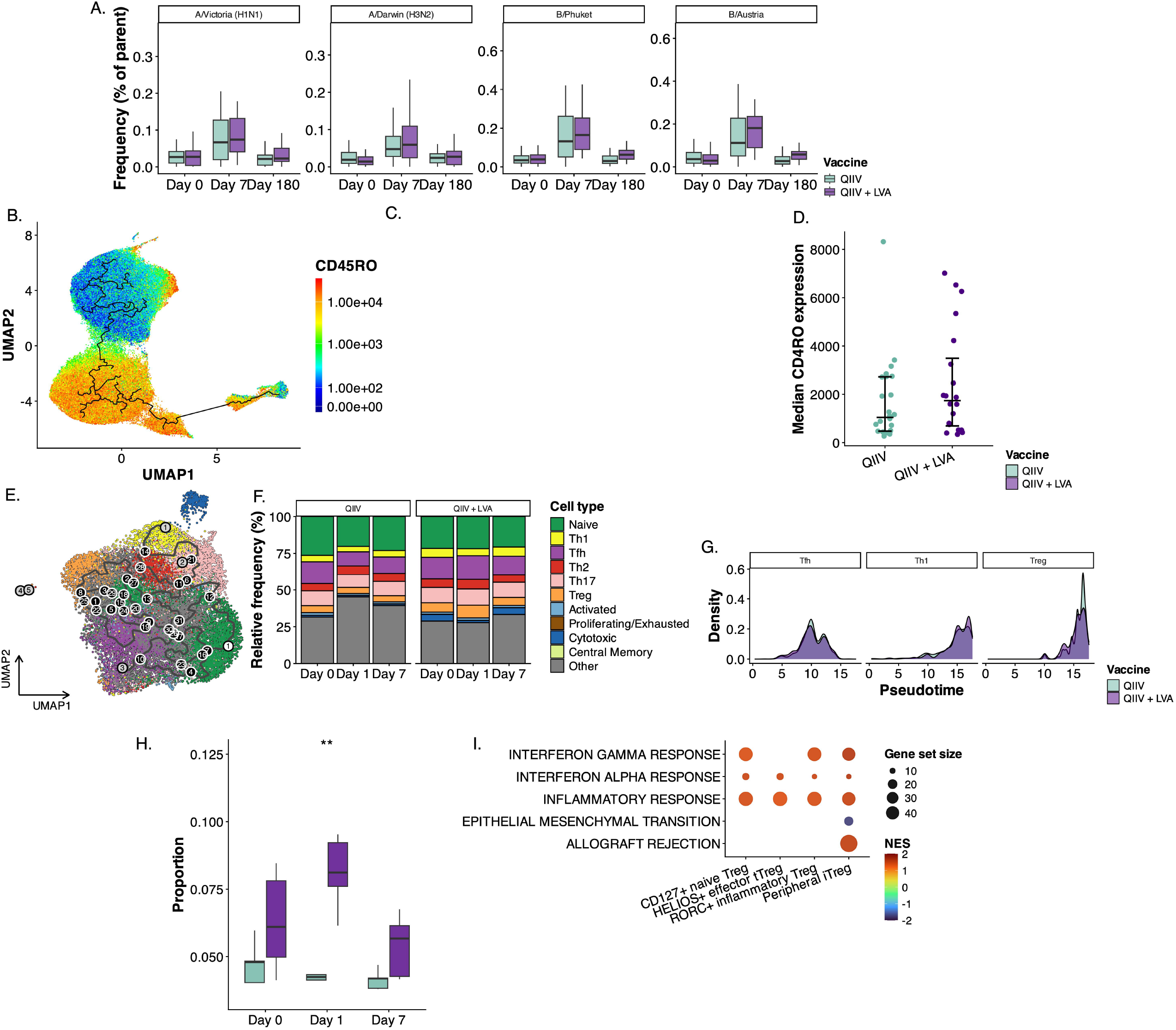
CD4+ T cell dynamics and differentiation trajectories following QIIV + LVA. (A) Antigen-specific CD4^+^ T cell responses measured by intracellular cytokine staining and flow cytometry following stimulation with influenza strains included in the vaccine (A/Victoria H1N1, A/Darwin H3N2, B/Phuket, B/Austria). Frequencies of polypositive CD4+ T cells are shown at days 0, 7, and 180 post-vaccination. Both vaccines induce antigen-specific T cell responses across multiple strains with a peak at day 7. (B) UMAP visualization of gated CD4^+^ T cell (averaged across influenza-strain stimulation conditions; flow cytometry data) states colored by CD45RO marker intensity, with the pseudotime trajectory overlaid, illustrating differentiation trajectories from naïve to activated and memory-like states following vaccination. (C) Density distribution of gated CD4+ T cells (flow cytometry data) along pseudotime bins for each vaccine group. QIIV + LVA shows a shift toward more advanced differentiation states compared to QIIV alone. (D) Median + range of CD45RO marker intensity of gated CD4+ T cells per vaccine group (flow cytometry data). Increased CD45RO expression indicates acquisition of memory phenotypes, which was not significantly higher in the QIIV + LVA group. (E) UMAP representation of CD4+ T cell subsets (in single-cell RNA sequencing) across timepoints (days 0, 1, and 7) and vaccine conditions, showing the identification of naïve, effector (Th1, Th2, Th17), Tfh-like, regulatory T cells (Treg), cytotoxic, proliferating/exhausted, and memory populations, overlaid with pseudotime trajectory (numbers and line). (F) Relative frequencies of CD4+ T cell subsets over time. Vaccination induces limited remodeling of CD4+ T cell composition, with expansion of effector and Tfh-like populations, slightly more pronounced in the QIIV + LVA group. (G) Density distribution of specific CD4+ T cell subsets (Tfh, Th1, and Treg) along pseudotime, highlighting differential positioning of these subsets within the differentiation trajectory. (H) Proportions of total Treg over time. Vaccination alters the balance of Treg subsets, with significantly elevated levels observed early after QIIV + LVA immunization. (I) Gene set enrichment analysis (GSEA) across Treg subpopulations, including CD127+ naïve Tregs, HELIOS+ thymic-derived effector Tregs (tTreg), RORC+ inflammatory Tregs, and peripheral induced Tregs (iTreg), of Hallmark pathways at day 1 post-vaccination between vaccine groups. Normalized enrichment scores (NES) for significant pathways are shown and indicate enrichment for QIIV + LVA in red or enrichment in QIIV alone in blue. Dot size indicates the gene set size. Hallmark pathways associated with inflammation and interferon signaling are enriched during differentiation, with higher NES in the QIIV + LVA condition. Significance levels are indicated as * (p < 0.05), ** (p < 0.01), *** (p < 0.001), **** (p < 0.0001), unless otherwise specified. LVA: Litevax Adjuvant; NES: normalized enrichment score; QIIV: quadrivalent inactivated seasonal influenza vaccine;

Unsupervised analysis of gated CD4+ T cells (**Extended Figure 3A**) revealed a continuum of differentiation states captured by pseudotime inference (**Extended Data Figure 3C&D**). Mapping CD45RO expression onto this trajectory showed increasing marker intensity along pseudotime, with the highest expression observed in later pseudotimes (**Figure 3B**). As CD45RO is associated with antigen-experienced and memory T cells, this suggests that the inferred trajectory reflects progressive transitions towards more activated or memory-like states. Importantly, while overall CD45RO expression levels were comparable between vaccine groups (**Figure 3D**), cell density analysis across pseudotime bins demonstrated significant enrichment of CD4+ T cells in later pseudotime states in the QIIV□+□LVA group (bin 10-15: p <0.0001; bin 15-20: p <0.0001; bin 25-30: p <0.0001) (**Figure 3C**). These findings indicate that LVA does not alter CD45RO expression per se but is associated with a redistribution of CD4+ T cells along the inferred cellular continuum, with a greater proportion of cells in CD45RO-high states. Notably, pseudotime analysis enabled the detection of continuous shifts in cellular states that were not apparent from conventional analyses of total influenza-specific CD4+ T-cell frequencies.

Phenotypic classification based on CCR7 and CD45RO expression further confirmed the presence of naïve, central memory, effector memory, and terminal effector subsets (**Extended Data Figure 3E-H**), but did not reveal major differences in their overall frequencies between vaccine groups. The frequency of central memory CD4+ T cells was significantly higher in the QIIV + LVA group at day 7 (0.25±0.36% versus 0.13±0.26%; p = 0.009) and day 180 (0.26±0.37% versus 0.15±0.32%; p = 0.014). However, a similar difference was already present at baseline (0.26±0.35% versus 0.16±0.31%, p = 0.014), indicating that these observations likely reflect pre-existing variation rather than vaccine-induced expansion of central memory cells (**Extended Data Figure 3F**).

To further dissect the differentiation trajectories underlying these responses, we analyzed single-cell RNA sequencing data from CD4+ T cells. Unsupervised clustering identified diverse T helper subsets, including Th1, Tfh-like, Th2, Th17, and regulatory T cell (Treg) populations (**Figure 3E**). Relative abundance analyses did not reveal significant changes in the overall proportion of Th1 or Tfh-like cells following both QIIV and QIIV□+□LVA administration (**Figure 3F**; **Extended Data Figure 3I&J**), indicating that LVA does not markedly alter the composition of canonical T helper subsets.

Trajectory inference, however, revealed subtle (n.s.) but structured differences in the distribution of CD4+ T cells along differentiation pseudotime following LVA administration. Specifically, modest increases in the densities of Tfh-like cells were observed in early pseudotime bins, whereas Th1 cells exhibited higher density peaks in later pseudotime states in the QIIV□+□LVA group (**Figure 3G**). These findings suggest that, rather than reshaping overall subset frequencies, LVA is associated with a redistribution of CD4+ T cells along inferred pseudotime trajectories consistent with predicted progression towards T helper states.

Single-cell analyses revealed a significant increase in the proportion of regulatory T cells (Tregs) at day 1 following QIIV□+□LVA compared with QIIV alone (proportion: 0.022 vs. 0.004, p = 0.013) (**Figure 3H**).

To further characterize the functional properties of these cells, Tregs were subsetted and re-annotated, revealing distinct subpopulations, including peripheral and thymic-derived Tregs (**Extended Data Figure 3K**). Pathway enrichment analysis demonstrated enrichment of IFN*γ* responses in naïve Tregs (NES: 1.36, p = 0.022), RORC+ Tregs (NES: 1.42, p = 0.006) and peripheral Tregs (pTreg) (NES: 1.59 p = 0.006) in the QIIV□+□LVA group. Notably, only pTregs exhibited enrichment of the allograft rejection pathway, a gene set associated with antigen presentation and T cell activation (NES: 1.52, p = 0.004) (**Figure 3I**), suggesting a functionally active phenotype. These findings are consistent with the identification of CTLA4 as a receptor engaged by dendritic cell-derived signals. Together, the increased abundance and activation state of Tregs are consistent with the engagement of regulatory pathways alongside effector T cell priming following LVA administration.

Collectively, these data show that LVA does not increase the magnitude of influenza-specific CD4+ T cell responses but is associated with qualitative differences in CD4+ T cell states, including enrichment of CD45RO+ pseudotime states, subtle shifts towards Th1- and Tfh-like phenotypes, and early activation of regulatory T cell programs. These observations are consistent with an effect of LVA on the cellular composition and functional programming of the adaptive T cell response.

### LVA is associated with enhanced early IgG3 responses

Analysis of IgG subclass responses revealed similar kinetics of all IgG subclasses between the two vaccine groups (**Figure 4A**; **Extended Data Figure 4A**). No significant differences in the overall magnitude of total IgG responses or hemagglutination inhibition (HI) titers were observed between QIIV and QIIV□+□ LVA groups across time points, consistent with our previously published findings (16). In both groups, antibody titers increased following vaccination and subsequently stabilized, reflecting the expected kinetics of recall responses to seasonal influenza vaccination. Longitudinal modelling using estimated marginal means (EMMs), accounting for repeated measures and inter-individual variability, recapitulated these trajectories and showed strong overlap between groups at later time points (**Extended Data Figure 4B&C**). These findings prompted us to further investigate potential differences in early IgG subclass responses.

**Figure 4.**
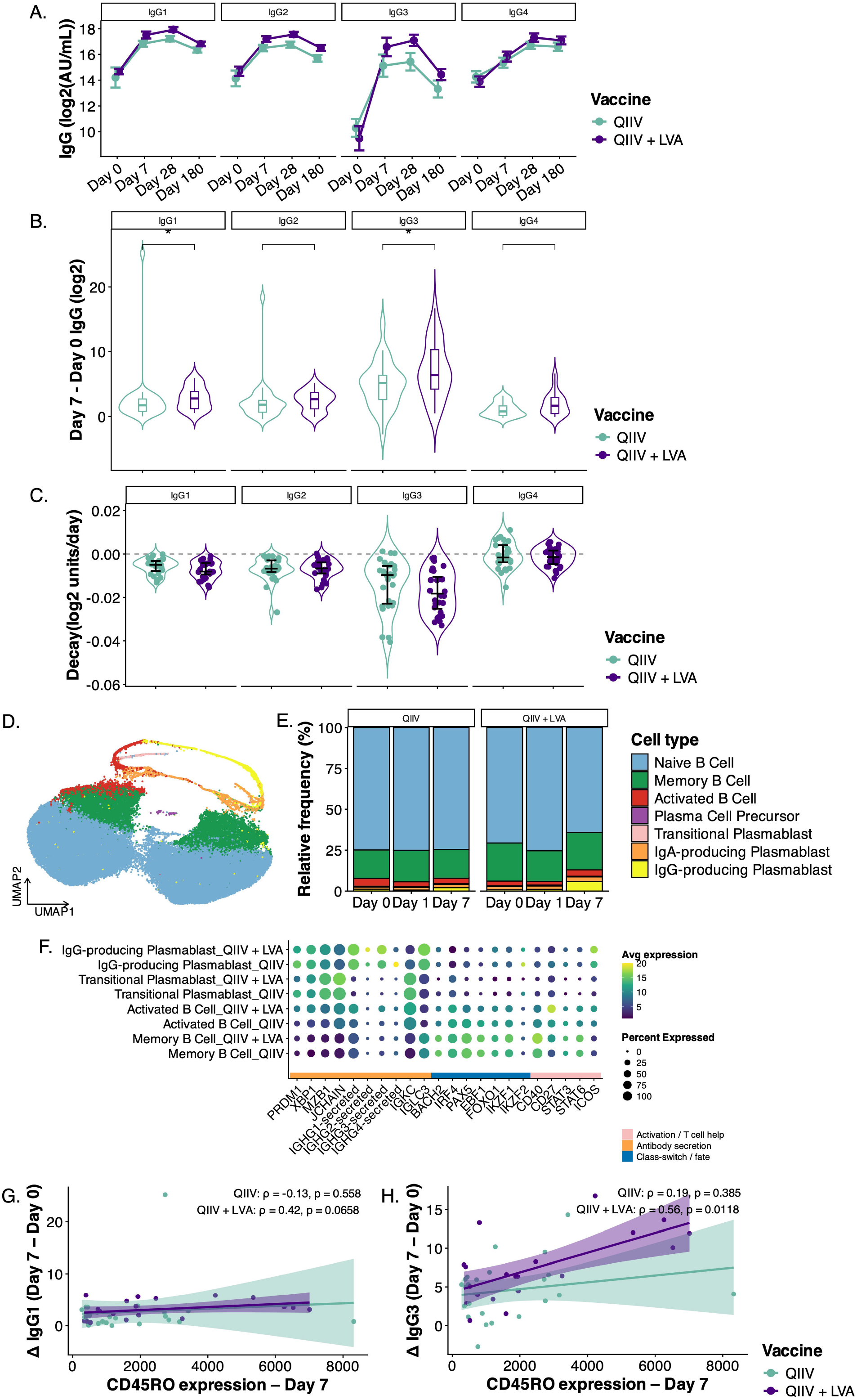
Humoral immune responses and B cell dynamics following QIIV + LVA. (A) Longitudinal serum IgG subclass responses (IgG1, IgG2, IgG3, and IgG4) following vaccination at days 0, 7, 28, and 180. Mean + SEM Antibody titers (log_2_ AU/mL) increase after vaccination, with a peak at day 28 but without significant differences between vaccine groups. (B) Violin- and boxplot indicating the slope (as a measure of increase) of IgG subclass levels from day 0 to day 7 (ΔIgG). Early antibody responses show subclass-specific differences, with significantly stronger increases of IgG1 and IgG3 in the QIIV + LVA group. (C) Violin- and whiskers plot of estimated decay rates of IgG subclasses over time (from day 28 until day 180), expressed as log_2_ units per day. These analyses reveal no differences in antibody durability between vaccine formulations. (D) UMAP embedding of single-cell RNA-sequencing data of B cell populations. Major subsets include naïve B cells, memory B cells, activated B cells, plasmablasts, plasma cell precursors, and immunoglobulin class–specific plasmablasts (IgA- and IgG-producing). (E) Relative frequencies of B cell subsets across timepoints (days 0, 1, and 7) and vaccine groups. Vaccination induces expansion of activated B cells and plasmablast populations, with stronger responses in the QIIV + LVA condition. (F) Dot plot of gene expression across activated B cell subsets. Markers related to activation and T cell help (CD40, CD27, ICOS, STAT3/6), antibody secretion (PRDM1, XBP1, MZB1, JCHAIN), class-switch recombination and cell fate (BACH2, IRF4, EBF1, IKZF1/2) define functional states of B cell differentiation. Dot size represents the percentage of expressing cells and colour indicates average expression. (G) Correlation between CD4+ T cell activation (CD45RO marker intensity at day 7) and early IgG1 response (ΔIgG1, day 7 minus day 0). A positive association (n.s.) is observed in the QIIV + LVA group but not in QIIV alone. (H) Correlation between CD4+ T cell activation (CD45RO marker intensity at day 7) and early IgG3 response. A stronger and significant positive correlation is observed in the QIIV + LVA group, suggesting enhanced T cell help promotes class-switched antibody responses. Significance levels are indicated as * (p < 0.05), ** (p < 0.01), *** (p < 0.001), **** (p < 0.0001), unless otherwise specified. LVA: Litevax Adjuvant; QIIV: quadrivalent inactivated seasonal influenza vaccine;

To further characterize early humoral responses, we assessed the rate of increase in IgG subclass levels between day 0 and day 7. This analysis revealed significantly steeper increases in both IgG1 (log2-fold change: 2.84±1.76 vs. 2.62±4.56, p = 0.048) and IgG3 (log2-fold change: 7.10±4.30 vs. 4.84±3.56, p = 0.046) in the QIIV□+□LVA group compared with QIIV alone (**Figure 4B**), indicating accelerated early antibody responses. Given the association between IgG3 class switching and inflammatory cytokine and interferon signaling, these findings are consistent with the systemic interferon response induced by LVA. In contrast, decay rates between day 28 and day 180 were not significantly different between groups across all IgG subclasses (**Figure 4C**), indicating that LVA mainly affects early response kinetics.

To investigate the cellular basis of these observations, we analyzed B cell populations using single-cell RNA sequencing. Unsupervised clustering identified the major B cell subsets, including naïve B cells, memory B cells, activated B cells, and plasmablasts (**Figure 4D**). Following vaccination, the QIIV□+□LVA group showed a higher proportion of IgG-producing plasmablasts at day 7 compared with the QIIV group (proportion: 0.059±0.022 vs. 0.021±0.131, p = 0.032) (**Figure 4E**; **Extended Data Figure 4D**).

Although the frequencies of memory B cells did not differ significantly between vaccine groups (**Extended Data Figure 4E**), transcriptional profiling of memory B cells revealed increased expression of genes associated with B cell activation and T cell help, including CD40, ICOS, and STAT3, in QIIV□+□LVA group (**Figure 4F**). In parallel, increased expression of immunoglobulin heavy chain transcripts, particularly IGHG3, was observed following QIIV□+□LVA administration, supporting the preferential induction of IgG3-producing cells. Together, these transcriptional signatures suggest enhanced responsiveness to T cell-derived signals.

To explore the relationship between CD4+ T cell activation and early antibody responses, we performed correlation analyses of CD4+ T cell phenotypes with IgG subclass responses. CD45RO expression at day 7 positively correlated with the magnitude of the IgG3 increase between day 0 and day 7 in the QIIV□+□LVA group (Spearman ρ = 0.56, p = 0.012) (**Figure 4H**), whereas the association with IgG1 responses did not reach statistical significance (Spearman ρ = 0.42, p = 0.06) (**Figure 4G**). In contrast, frequencies of effector memory CD4+ T cells were not associated with the slopes of either IgG1 or IgG3 responses (**Extended Data Figure 4F&G**), suggesting an association between CD45RO expression and early humoral responses that was not observed for the frequency of memory T-cell subsets.

Finally, we examined potential regulatory influences by assessing associations with CTLA4 expression. CTLA4 expression at day 1 negatively correlated with IgG1 levels at day 180 in the QIIV group (Spearman ρ = -0.54, p = 0.003) whereas no such association was observed in the QIIV□+□LVA group (Spearman ρ = 0.15, p = 0.45) (**Extended Data Figure 4H**). No significant associations between CTLA4 expression and IgG3 responses were detected in either vaccine group (**Extended Data Figure 4I**). These findings are consistent with a potential relationship between CTLA4-associated regulatory programs and long-term IgG1 responses in the QIIV group; however, this association was not observed following LVA administration.

Collectively, these data indicate that LVA does not substantially alter the overall magnitude and persistence of humoral immunity but is associated with accelerated early antibody production and enhances early IgG3 responses. These effects coincide with changes in innate immune activation, CD4+ T cell differentiation, and B cell activation, suggesting that LVA influences the qualitative features of the vaccine-induced humoral response.

### LVA is associated with reduced age-related interferon signatures and enhanced early IgG3 responses in older adults

To investigate the impact of age on vaccine-induced immune responses, participants were stratified into younger (18–50 years) and older (60 years or older) adults and analyzed at baseline and following vaccination. At baseline, older adults exhibited significant differences in immune cell composition, with altered frequencies of several circulating immune cell subsets, including Th1 (0.025±0.026% versus. 0.048±0.037%, p = 0.0004), and central memory CD4+ cells (0.236±0.345% versus 0.165±0.318%, p = 0.034), compared with younger adults (**Figure 5A**). Consistent with these observations, baseline antibody titers were lower in older adults across all IgG subclasses (p = 0.024; p = 0.015; p = 0.002; p = 0.006 for IgG1, IgG2, IgG3, and IgG4, respectively) (**Figure 5B**), showing age-related differences in both cellular and humoral immune status prior to vaccination. At the transcriptional level, baseline gene expression profiles in older adults were enriched for genes associated with immune regulation and cellular exhaustion (**Extended Data Figure 5A**), indicating age-associated differences in baseline transcriptional profiles.

**Figure 5.**
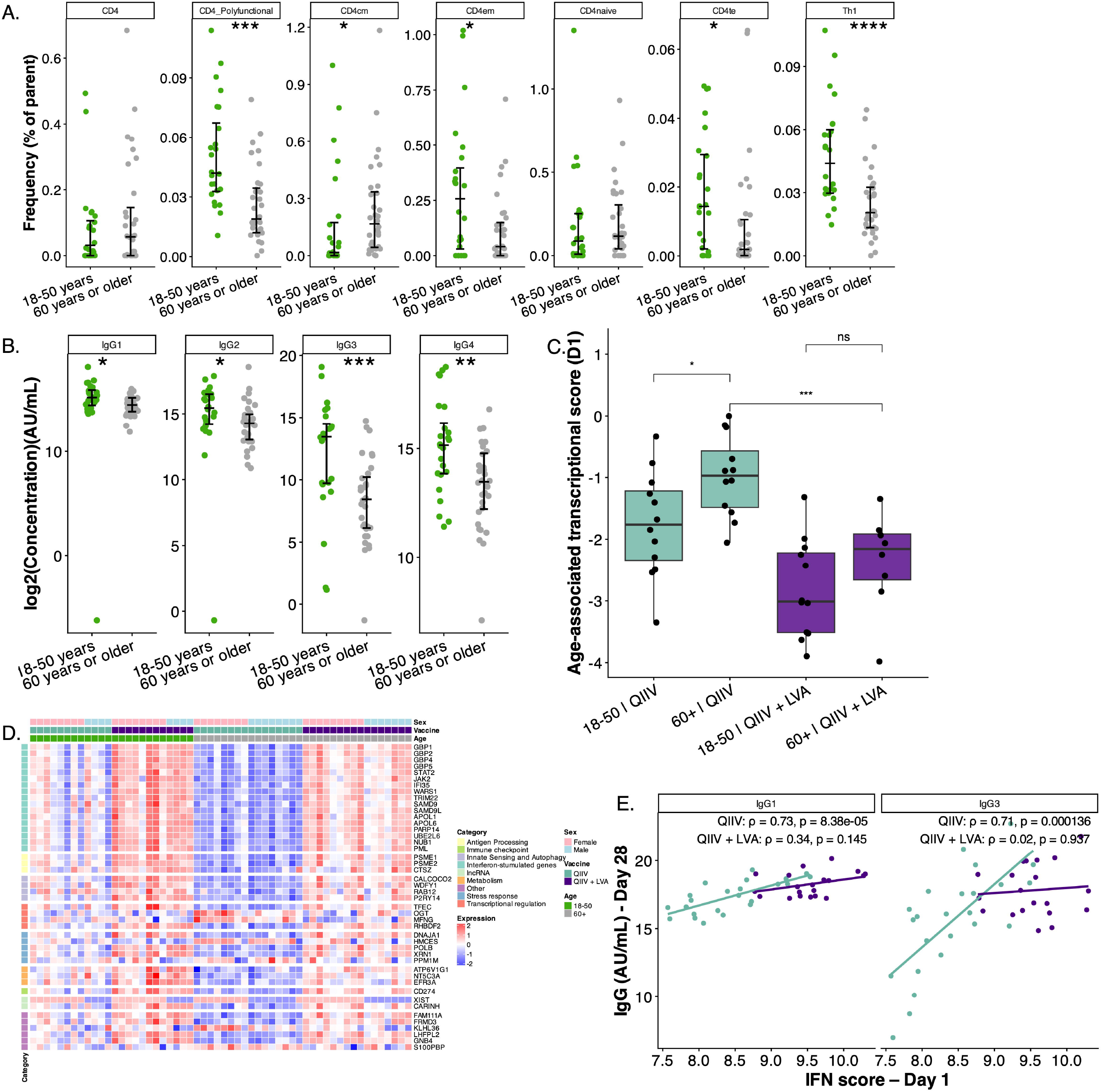
Interferon-associated modulation of vaccine responses in older adults. (A) Frequencies of CD4+ T cell subsets (from flow cytometry data and manual gating) stratified by age group at baseline (day 0). Subsets include total CD4+ T cells, polyfunctional CD4+ T cells, central memory (CD4cm), effector memory (CD4em), naïve (CD4naive), terminal effector (CD4te), and Th1 cells. Differences between younger and older individuals indicate age-dependent cellular immune responses to vaccination. (B) Mean + SEM serum IgG subclass responses (IgG1, IgG2, IgG3, and IgG4) at baseline (day 0), stratified by age and vaccine group. Antibody concentrations are reduced in older individuals. (C) Boxplot of age-associated transcriptional score at day 1 stratified by age- and vaccine group. The score was calculated for each sample as the average expression of genes upregulated in older adults minus the average expression of genes downregulated in older adults. As such, higher scores indicate a transcriptional profile more closely resembling that of older individuals at day 1, whereas lower scores reflect a profile more similar to younger individuals. Scores were not significantly different between YA and OA at day 1 after QIIV + LVA but were significantly higher in OA receiving QIIV, indicating that LVA transcriptionally regulates genes impacted in older adults. (D) Heatmap of genes part of the Hallmark IFN*γ* response pathway, stratified by sex, vaccine group, and age category. Genes are grouped by functional categories, including interferon-stimulated genes, antigen processing, innate sensing and autophagy, immune checkpoints, metabolism, and transcriptional regulation. Color scale indicates normalized expression (z-score). Distinct transcriptional signatures are observed across sex (female vs male), vaccine (QIIV vs QIIV + LVA), and age groups (18–50 vs ≥60 years), highlighting context-specific immune activation programs. (E) Correlation between age-associated transcriptional score at day 1 and antibody responses (IgG1 and IgG3) at day 28. A positive association is observed in the QIIV group, whereas this relationship is diminished in the QIIV + LVA group, suggesting altered coupling between early innate responses and downstream humoral immunity. Significance levels are indicated as * (p < 0.05), ** (p < 0.01), *** (p < 0.001), **** (p < 0.0001), unless otherwise specified. CD4cm: CD4+ central memory T cell; CD4em: CD4+ effector memory T cell; CD4te: CD4+ terminal effector T cell; LVA: Litevax Adjuvant; OA: older adults; QIIV: quadrivalent inactivated seasonal influenza vaccine; YA: younger adults.

To determine how these differences evolved following vaccination, we quantified age-associated transcriptional responses on day 1 by deriving a composite gene expression score based on genes differentially expressed between older and younger adults (**Extended Data Figure 5B**). Specifically, the score for each sample was calculated as the average expression of genes upregulated in older adults minus the average expression of genes downregulated in older adults. Higher scores therefore indicate a transcriptional profile more closely resembling that observed in older adults at day 1, whereas lower scores reflect a profile more similar to that of younger adults. Notably, many of the genes contributing most strongly to this score were interferon-stimulated genes, indicating that the metric captures an interferon-enriched transcriptional program associated with age-related differences in the day 1 response.

Older adults receiving QIIV alone exhibited significantly higher day 1 scores compared to younger adults, as well as compared to older adults receiving QIIV□+□LVA (**Figure 5C**). In contrast, no significant differences were observed between older and younger adults in the QIIV□+□LVA group, suggesting that LVA attenuates age-associated transcriptional differences at this early time point. Visualization of IFN*γ* response genes further demonstrated segregation by both age and vaccine group, with older adults in the QIIV group displaying lower expression of interferon-stimulated genes than younger adults (**Figure 5D**). This age-related transcriptional pattern was less pronounced following QIIV□+□LVA administration, where expression of interferon-associated genes was more comparable between age groups. Together, these findings indicate that age-associated differences in the early transcriptional response are largely driven by coordinated variation in interferon-related pathways and suggest that LVA modifies the magnitude and distribution of these responses across individuals.

We next assessed whether these early transcriptional responses were associated with downstream antibody outcomes. In the QIIV group, the age-associated transcriptional score positively correlated with both IgG1 (Spearman ρ = 0.73, p < 0.0001) and IgG3 titers at day 28 (Spearman ρ = 0.71, p = 0.0001) (**Figure 5E**), indicating that individuals with stronger day 1 transcriptional responses subsequently mounted higher antibody levels. These associations remained significant when analyzed separately within younger and older adults (**Extended Data Figure 5D**), suggesting that they reflect inter-individual variation rather than age stratification alone. In contrast, no significant correlations were observed in the QIIV□+□LVA group (**Figure 5E**; **Extended Data Figure 5D**), indicating that LVA alters the relationship between early transcriptional responses and subsequent antibody magnitude.

Consistent with this interpretation, no association was observed between the age-associated transcriptional score and CD45RO expression in CD4+ T cells at day 7 (**Extended Data Figure 5C**), suggesting that this transcriptional program is not directly associated with the distribution of CD4+ T cell states captured by CD45RO expression.

Collectively, these findings show that older adults exhibit a distinct interferon-enriched transcriptional response following seasonal influenza vaccination and that, in the QIIV group, this response is associated with subsequent antibody production. In contrast, these age-associated transcriptional signatures were less apparent in the QIIV + LVA group, and no significant association was observed between early transcriptional signatures and later humoral responses. Together, these results indicate that the relationships between age, early transcriptional responses, and antibody production differ between QIIV + LVA and QIIV.

## Discussion

In this study, we performed a longitudinal multi-omics analysis to define the mechanisms underlying the mode of action of LVA when administered with a quadrivalent inactivated seasonal influenza vaccine. We found that LVA induces a rapid and transient innate immune response characterized by a dominant interferon-driven transcriptional program, primarily within monocytes. This early innate activation was associated with enhanced antigen presentation by DCs and increased cell-cell communication with CD4+ T cells. At the adaptive level, LVA promoted a redistribution of CD4+ T cells towards more differentiated CD45RO+ memory states without increasing the overall frequency of antigen-specific cells. These changes were accompanied by accelerated early B cell activation, increased plasmablast responses, and enhanced early IgG3 antibody responses. Importantly, these effects were particularly relevant in older adults, in whom LVA attenuated age-associated transcriptional differences and reduced the relationship between pre-existing immune states and vaccine responsiveness. Collectively, our findings identify interferon signaling as a central axis linking innate and adaptive immunity following LVA-adjuvanted vaccination.

The observation that LVA induces a rapid and coordinated interferon-dominated response is consistent with systems vaccinology studies of influenza vaccines, that identified early interferon-stimulated gene signatures as predictors of antibody responses (18, 19). However, whereas previous studies largely established interferon responses as correlates of immunogenicity, our findings suggest that, in the context of LVA, interferon signaling may play a more prominent role in coordinating immune responses across multiple cellular compartments. This distinguishes LVA from an adjuvant such as MF59, which predominantly induces chemokine-driven recruitment of innate immune cells without a similarly centralized interferon program (20).

At the cellular level, monocytes exhibited the strongest enrichment of interferon-associated pathways, consistent with studies showing that interferon-driven activation of monocytes and DCs contributes to effective vaccine-induced immunity (21). In MF59-adjuvanted vaccines, monocytes are rapidly recruited and differentiate into dendritic-like cells that facilitate antigen transport and presentation (22, 23). In contrast, our findings suggest that LVA may preferentially promote functional reprogramming of monocytes towards an interferon-responsive state rather than inducing major shifts in cellular composition. The interferon-dominated response induced by LVA was accompanied by qualitative enhancement of antigen-presenting cell function. Although broad upregulation of MHC class II genes was not observed, dendritic cells displayed increased expression of the antigen-processing component CD74 and enhanced predicted co-stimulatory interactions with T cells. These findings are consistent with the concept that effective adjuvants activate dendritic cells to facilitate T cell priming, as has been described for AS01 (24, 25). Notably, LVA also induced classical inflammatory pathways, including TNFα and IL-6 signaling, indicating that its activity extends beyond interferon induction alone.

Within the T cell compartment, LVA did not increase the magnitude of influenza-specific CD4+ T cell responses. Instead, it was associated with a redistribution of CD4+ cells towards more differentiated CD45RO+ states along pseudotime trajectories. These observations support the concept that adjuvants shape the quality and differentiation of T cell responses rather than simply their magnitude (26). Consistent with the established role of interferon signaling in promoting Th1 and Tfh differentiation, these findings suggest that LVA enhances functional maturation of helper T cell responses. Compared with MF59, which has primarily been associated with enhanced humoral immunity and more limited effects on T cell polarization (20, 27, 28), LVA may exert a broader influence on T cell differentiation pathways.

These coordinated innate and T cell responses were mirrored in the humoral compartment. LVA accelerated plasmablast responses and selectively enhanced early IgG3 class switching. This observation is consistent with evidence implicating interferons and inflammatory cytokines in promoting switching towards cytophilic IgG subclasses, including IgG3 (29). In contrast to MF59, which has been shown to increase overall antibody titers and augment antibody effector functions (20, 30), LVA did not increase total IgG responses. Instead, it appeared to influence the kinetics and subclass distribution of the antibody response, suggesting that its effects are predominantly qualitative rather than quantitative. Our findings extend previous *in vitro* observations showing that CMS-based adjuvants activate antigen-presenting cells and promote T helper polarization (15). Prior studies have demonstrated that CMS can activate dendritic cells through TLR2 and TLR4-dependent pathways, resulting in cytokine production and enhanced antigen presentation (15, 16). Here, we provide evidence that these translate to humans *in vivo* and are associated with coordinated immune responses spanning innate and adaptive compartments.

The inclusion of both younger and older adults enabled direct assessment of LVA activity in the context of immunosenescence. Consistent with previous reports, older adults exhibited baseline alterations in immune composition and function, including reduced antibody titers and transcriptional signatures associated with immune dysregulation and exhaustion (31). These findings align with extensive evidence demonstrating that aging is associated with impaired adaptive immunity, reduced B cell function, and diminished vaccine responsiveness (7, 31, 32).

Dysregulation of innate immune signaling, including impaired interferon responses, represents a key feature of immunosenescence (8, 32). Reduced interferon responses have been linked to impaired antigen-presenting cell activation and diminished adaptive immune priming, highlighting its importance as an early determinant of vaccine-induced immunity (32). In line with these observations, previous studies have demonstrated that early interferon-related transcriptional signatures predict vaccine responsiveness across diverse populations, including older adults (19).

In the absence of adjuvant, early interferon-associated transcriptional responses correlated with subsequent antibody responses, suggesting that inter-individual variability in innate immune activation contributes to heterogeneity in vaccine responsiveness. In contrast, these relationships were not observed following LVA administration. This finding suggests that the relationship between pre-existing immune states and vaccine-induced responses may differ following LVA administration. Notably, LVA also attenuated age-associated differences in early transcriptional responses, resulting in more comparable interferon-associated profiles between younger and older adults. Given that aging has been associated with impaired interferon production and signaling, enhancement of this pathway may help restore effective coordination between innate and adaptive immunity in older adults.

The enhanced early IgG3 response observed following LVA administration may be particularly relevant in this context. IgG3 exhibits potent effector functions, including complement activation and high-affinity Fc receptor engagement, and may therefore contribute to protective immunity even in the absence of higher antibody titers.

This study has several strengths. First, the integration of bulk transcriptomics, single-cell RNA sequencing, flow cytometry, and serology provides a comprehensive, systems-level view of vaccine-induced immunity. Second, the longitudinal design captures both early innate responses and downstream adaptive outcomes, enabling mechanistic connections between these processes. Third, the inclusion of both younger and older adults allows direct assessment of age-related differences and enhances the translational relevance of the findings. Finally, the combined use of computational and experimental approaches to assess cell–cell communication provides novel insights into how adjuvants modulate immune network interactions.

Several limitations should also be acknowledged. First, the study is observational, and although strong associations were identified, causal relationships, particularly regarding the central role of interferon signaling, cannot be definitively established. Experimental perturbation studies will be required to validate these mechanisms. Second, older adults were not included in the single-cell RNA sequencing analysis. This decision was made to focus the mechanistic analyses on younger adults, in whom vaccine-induced immune responses were expected to be more robust and homogeneous, thereby facilitating the identification of cellular and molecular pathways associated with LVA activity. Nevertheless, this approach precludes direct assessment of age-specific cellular dynamics and transcriptional responses at single-cell resolution in the population most relevant to immunosenescence. Furthermore, the single-cell analyses were performed in a small, selected subset of participants (n = 3 per group) and should therefore be considered exploratory and hypothesis-generating. As a result, the observed differences in cellular states, inferred trajectories, and predicted cell-cell communication networks require validation in larger cohorts. Computational approaches such as pseudotime inference and CellChat analysis generate statistical predictions regarding cellular state transitions and intercellular communication but do not provide direct functional evidence of differentiation processes or signaling interactions. These findings should therefore be interpreted with appropriate caution until confirmed through independent datasets and functional experiments. Third, our analyses focused on peripheral blood responses and may not fully capture immune processes at the site of vaccination, within lymphoid tissues where critical priming events take place, or at mucosal sites in the lung relevant for protection against influenza. Finally, validation in independent cohorts will be important to confirm the robustness and reproducibility of these findings. Moreover, in the absence of established correlates of protection and vaccine efficacy endpoints, it remains unclear whether the immunological changes associated with LVA translate into improved clinical protection against influenza.

In conclusion, our study provides novel insights into the immunological effects of LVA when administered with a seasonal influenza vaccine. LVA was associated with a pronounced interferon response, altered innate immune signatures, qualitative changes in adaptive immune cell states, and shifts in antibody isotype profiles. Together, these findings suggest that LVA influences multiple components of the vaccine-induced immune response, particularly its qualitative characteristics rather than its overall magnitude. In older adults, LVA was associated with differences in age-related transcriptional signatures, warranting further investigation into the potential impact of interferon-inducing adjuvants on vaccine responses across the lifespan. While the underlying mechanisms and clinical implications require further validation, our results provide mechanistic insights into the immunomodulatory effects of interferon-inducing adjuvants and support their further investigation as strategies to optimize vaccine responses, particularly in populations with diminished vaccine responsiveness.

## Methods

### Study design and participants

This study was a randomized, double-blind, active-controlled phase 1b trial conducted at a single center, the Center for Vaccinology (CEVAC, Ghent University Hospital and Ghent University in Ghent, Belgium), between January 22 and October 10, 2024. The study enrolled 84 healthy male and female adults, including 36 participants aged between 18 and 50 years and 48 participants over 60 years. Details on the study design have been previously published (17). All study procedures were conducted in accordance with the International Council for Harmonisation (ICH) guidelines for Good Clinical Practice (GCP). The study documents were approved by an independent Ethics Committee and the Belgian Federal Agency for Medicines and Health Products (FAMHP) (EudraCT number: 2023-508230-33-00, NCT06294262). Written informed consent was obtained from all participants prior to any study-related procedures.

### Study vaccine

The adjuvant formulation consisted of CMS combined with a squalane-in-water emulsion. This fully synthetic, sterile, aqueous, ready-for-use product is characterized by its physical and chemical stability. In this study, LVA containing 0.5mg or 1mg CMS was mixed with VaxigripTetra (QIIV) (Sanofi Pasteur, Lyon, France), a licensed quadrivalent seasonal inactivated split-virion influenza vaccine, immediately prior to intramuscular administration. Each vaccine dose contained 15 µg of HA per strain and was administered either alone or combined with 0.5mg or 1mg of CMS. The 2023-2024 Northern Hemisphere formulation of QIIV, composed according to WHO’s recommendations for that season, was used in all study groups. The injection volume was 0.5 mL for QIIV and 0.55 mL for QIIV combined with LVA. Only blood samples from participants receiving QIIV alone or QIIV + 1 mg LVA were analyzed here, resulting in a total of 56 participants included in this analysis.

### Sample collection

Blood samples were collected from all participants at baseline (day 0) and on days 1, 7, 28 and 180. Influenza-like illnesses were monitored continuously during the study using nasal self-tests in symptomatic participants to assess the potential impact of intercurrent infections on observed immune responses. Five mL of blood was collected by venous puncture in serum separation blood collection tubes (Becton Dickinson Vacutainer tubes). Serum was collected after centrifugation for 10 minutes at 1300-2000g and frozen at 400µL per aliquot. Fifty mL of blood was collected by venous puncture in heparin-coated blood collection tubes (Becton Dickinson Vacutainer tubes of 10 mL coated with lithium heparin) from all participants at baseline (day 0) and at day 1, day 7, day 28 and day 180. After approximately 1:2 dilution in Hanks’ buffered salt solution (HBSS), PBMCs were isolated by density gradient centrifugation (Lymphoprep™), washed twice in HBSS, suspended in freezing solution (10% dimethyl sulfoxide/90% fetal bovine serum v/v), frozen at a concentration of 5 up to 20 million cells/mL, and stored in liquid nitrogen until use in the respective assays.

### Intracellular cytokine staining and flow cytometry

PBMCs from all participants collected at baseline (day 0), day 7 and day 180 were thawed and suspended in RPMI 1640 medium supplemented with Minimum Essential Medium non-essential amino acids, L-glutamine, penicillin/streptomycin, sodium pyruvate, 2-mercapto-ethanol (all from Invitrogen), and heat-inactivated fetal bovine serum (Seradigm). Then, PBMCs were incubated in vitro with peptide pools of the relevant vaccine antigens (HA/H3N2/Darwin/2021, HA/H1N1/Victoria/2022, HA/H0N0/Austria/2021 and HA/H0N0/Phuket/2013 (PepMix™, JPT Peptide Technologies GmbH, Germany) or left unstimulated (background condition with 0.32% DMSO) in the presence of costimulatory antibodies to CD28 and CD49d. Brefeldin A, a protein transport inhibitor, was added after 2 hours for subsequent overnight culture. The following day, cells were stained using fluorochrome-conjugated antibodies to phenotypic markers (CD3, CD4, CD8, CD45RO, CCR7), activation (CD40L, CD137, CD38, HLA-DR) and cytokine (IFN*γ*, IL2, IL13, IL17a TNF-α, and Granzyme B) markers. The stained cells were subsequently analyzed using flow cytometry.

### Immunoglobulin isotype analysis

Total IgG, and all 4 IgG subtypes were measured in all serum samples collected at baseline and 7-, 28-, and 180-days post-vaccination using the Meso Scale Discovery platform (MSD, Rockville, MD, USA) according to the manufacturer’s instructions. The MSD Flu-4 PLEX plates were custom-made plates, coated with the four antigens present in the 2023-2024 seasonal influenza vaccine: Flu A/Wisconsin/588/2019 H1, Flu B/Austria1359417/2021, Flu A/Darwin/6/2021 H3, and Flu B/Phuket/3073/2013. Briefly, frozen serum samples were thawed and analyzed in two dilutions. First, the MSD plate (Flu-4 PLEX 96-well 10-Spot plate, MSD) was blocked with Blocker A solution (150 µL/well) for at least 30 minutes, shaking at 700 revolutions per minute (rpm) at room temperature.

After incubation, the Blocker A solution was discarded, and the plate was washed three times using 150 µL/well of the 1X Wash Buffer. Next, 50 µL of either the standards, controls, or samples was added to the plate. For detection conditions IgG, IgG1, IgG2, and IgG3, the top of the curve for the standard dilution series was made by making a 10-fold dilution of Reference Standard 1 (Lot. A0080286). For IgG4 detection, undiluted Reference Standard 1 was used. The following calibrator dilutions (CAL02-CAL07) were prepared by performing a four-fold dilution series. All dilutions were made using Diluent 100. The final calibrator contained only Diluent 100 and was used as a blank. Serology Control 1.1 (Lot. A00C0825), 1.2 (Lot. A00C0826), and 1.3 (Lot. A00C0827) were supplied pre-diluted (1/5000) at the working concentration and were used for detection conditions IgG, IgG1, IgG2, and IgG3. These controls were replaced by three serum samples for the IgG4 analysis. The standard series and controls were added in duplicate. Depending on the detection condition, two different dilutions were tested to ensure that the unknown concentrations fell within detection ranges for at least one dilution. After standard, control, and sample addition, the plate was incubated for two hours at 700 rpm at room temperature. Before the end of incubation, 50 µL of the detection antibody was added (MSD GOLD SULFO-TAG Anti-Human IgG Antibody or the SULFO-TAG Anti-Human Antibody of the IgG subclass of interest) and incubated for 1 hour at 700 rpm at room temperature. After the final incubation step, the plate was washed three times using 150 µL/well of the 1X Wash buffer, and 150 µL/well of the MSD GOLD read buffer B was added to the plate. The readout of the plate happened within 15 minutes after the addition of the read buffer. Plates were analyzed using the MESO QuickPlex 120MM instrument and the Methodical Minds software (v2.1.3).

### Bulk RNA sequencing

Blood samples collected at baseline, on days 1 and 7 were collected in PAXgene tubes® (Qiagen, Hilden, Germany) maintained at room temperature for 2-72 hours, transferred to -20°C for short-term storage, and subsequently stored at -80°C until RNA extraction in batch. RNA was extracted using the Qiagen PAXgene blood RNA kit (IVD) according to the manufacturer’s instructions. RNA was stored in molecular grade water at -80°C until mRNA library preparation. Before starting library preparations, RNA yield and integrity were verified by microelectrophoresis using the Agilent Fragment Analyzer High Sensitivity kit in the Bioanalyzer 2100 system (Agilent, CA, USA). The cDNA libraries were constructed using the QuantSeq 3’ mRNASeq Library Prep Kit for Illumina FWD (Lexogen GmbH, Austria), following the manufacturer’s protocol. The final pool of libraries was subjected to a single-end sequencing (75 bp) with a sequencing read depth of 6 million reads.

### Single Cell RNA sequencing

A subset of six younger adults (three in each vaccine group) was selected based on their overall reactogenicity profile (CRP concentrations, white blood cell differentiation, and number and intensity of adverse events) as a marker of inflammation and innate immune activation. PBMCs collected at baseline (day 0), day 1 and day 7 were thawed, subjected to red blood cells lysis, and suspended in RPMI 1640 medium supplemented with Minimum Essential Medium non-essential amino acids, L-glutamine, penicillin/streptomycin, sodium pyruvate, 2-mercapto-ethanol (all from Invitrogen), and heat-inactivated fetal bovine serum (Seradigm). A targeted transcriptomics approach using the BD Rhapsody Express system (BD Biosciences) was performed. Samples were counted and resuspended in cold BD Sample Buffer (BD Biosciences) to achieve approximately 194,000 cells in 620 µL. Single cells were isolated using Single Cell Capture and cDNA Synthesis with the BD Rhapsody Express Single-Cell Analysis System following the manufacturer’s protocol (BD Biosciences). After priming the nanowell cartridges, the sample was loaded onto one BD Rhapsody cartridge and incubated at room temperature. Cell Capture Beads (BD Biosciences) were prepared and then loaded onto the cartridge and incubated. According to the manufacturer’s protocol, cartridges were washed, cells were lysed, and Cell Capture Beads were retrieved and washed prior to performing reverse transcription and treatment with Exonuclease I. cDNA Libraries were prepared using mRNA Targeted Library Preparation with the BD Rhapsody Targeted mRNA protocol (BD Biosciences). In brief, cDNA underwent targeted amplification using the Human Immune Response Panel primers and a custom supplemental panel via nested PCR (2×10 cycles) with Ampure XP bead cleaniup between amplifications. PCR products were then purified using Ampure XP beads. Quality and quantity of PCR products were determined by using an Agilent Fragment Analyzer High Sensitivity kit (Agilent). Targeted mRNA product was diluted to 2.7 ng/µL to prepare final libraries. Final libraries were indexed using PCR (6 cycles). Index PCR products were purified using Ampure XP beads. Quality of final libraries was assessed by using Agilent Fragment Analyzer High Sensitivity kit. Following library QC according to Illumina’s Sequencing Library qPCR Quantification Guide, libraries were pooled equimolarly based on qPCR quantification. The pooled library was loaded onto the AVITI sequencer at 9–12 pM and sequenced using an 8-0-51-71 run configuration (i1-i2-r1-r2; PE75). Libraries were multiplexed and sequenced to a total read depth of 6000 reads per cell.

### Bioinformatic analysis

#### Flow cytometry analysis

The samples were analyzed by flow cytometry (BD LSR Fortessa X-20, FlowJo v9.9.6). Influenza-specific CD4+ and CD8+ T cells were determined as CD3+CD4+ and CD3+CD8+ events expressing one marker or a combination of markers of CD40L, IFN-*γ*, IL-2, and TNF-α after *in vitro* stimulation with the vaccine antigen, from which the corresponding signal of the same sample obtained in the background condition was subtracted. The intracellular cytokine staining (ICS) results were reported as the frequencies (%) of influenza-specific CD4+ or CD8+ T cells per parent population. Results below 0.0001% after background subtraction were set at 0.0001%. All analyses have been done with ICS data multiplied by 10,000, resulting in the frequency of cells per million parent cells. Unsupervised analyses were performed in R using the unsupflowhelper package. Exploratory analyses indicated similar temporal patterns across strains, supporting the use of the mean response as a composite measure. Data was processed using the Phenograph clustering algorithm (as implemented in the R package Rphenograph (33)) to increase granularity of T cell subpopulations. Trajectory inference was conducted using the Cytotree R package (34). After subsetting CD4+ cells from total PBMCs, dimensionality reduction was applied via t-SNE, followed by trajectory construction using Cytotree’s tree-building algorithm. Pseudotime values were computed along the inferred trajectory, and cells were assigned to branches based on hierarchical clustering. Marker expression dynamics along pseudotime were assessed for key markers and cytokines.

#### Immunoglobulin isotype analysis

Raw data files (.txt files) were uploaded directly to the MSD Discovery Workbench software v4.0 (Meso Scale Discovery). MSD provided calibrator values for IgG antibodies specific for the antigens present in the Reference Standard. These values were applied to each IgG subclass and were used to create the calibration curve, where the mean signal of each calibrator level measured in duplicate was calculated and fitted to a four-parameter logistic (4PL) model using 1/Y2 weighting. The 4PL model estimated the upper asymptote, lower asymptote, slope, and midpoint of the sigmoidal response curve. The IgG4 assay used undiluted Reference Standard 1 as the top of the curve, so the calibrator values were multiplied by 10. A detection range was determined automatically, with the lower limit of detection set at a calculated concentration 2.5 standard deviations above the lowest point on the calibration curve, and the upper limit of detection set at the highest calibrator point. ECL signals of the samples were backfitted on the calibration curve, and the concentration was calculated in arbitrary units per mL (AU/mL). Samples were retested when values for both dilutions were above the detection range but not below. Samples with final values below the detection range were assigned half the value of the lowest measured concentration on the plate for that antigen. Samples above the detection range were assigned to have the highest measured concentration on the plate for that antigen. The final concentration was calculated as the average of both dilution values. For samples that were retested, only the two most recent concentrations within the detection range were used to determine the final concentration.

#### Bulk RNA sequencing analysis

Raw single-end reads were pre-processed for quality control. Sequencing quality was assessed before and after adapter trimming using FastQC (35). The Cutadapt software version 4.9 (36) was used to remove adapter sequences, trimming the 3′ ends with a mean quality score below 20 and discard reads shorter than 35 bp after trimming. After preprocessing, the high-quality reads were mapped onto the reference genome Homo sapiens (version GRCh38), and gene abundance was quantified using STAR (version 2.7.11a) (37) with default settings, 2-pass mapping and per gene read counts. These were done using the European Galaxy Servers (https://usegalaxy.eu/).

Principal component analysis (PCA) was performed to visualize data in a lower dimension and to investigate for confounders (batch of RNA extraction, age, biological sex and treatment group). Analysis of differentially expressed genes (DEGs) was performed using the R/Bioconductor package DESeq2 (38). Differential expression analysis (DEA) was performed for the full dataset to identify vaccine-specific changes at any time point. Additionally, DEA was done to identify DEGs at day 1 or day 7 compared to day 0 for each vaccine separately, and to identify DEGs for QIIV + LVA compared to QIIV alone at each day. All models included age and sex as confounders. Participant ID was not included as a blocking factor in the primary model; therefore, longitudinal DEG counts should be interpreted as descriptive of vaccine-associated transcriptional changes rather than as fully paired estimates. Wald test P⍰values were corrected for false⍰discovery rate using the Benjamini-Hochberg procedure, with statistical significance defined as an adjusted p⍰value ≤ 0.05 and a fold⍰change > 1. The obtained sets of DEGs were used for hierarchical clustering and visualized in heatmaps using pheatmap. Subsequently, all sets of DEGs were ranked and subjected to gene set enrichment analysis (GSEA) using the R/Bioconductor package clusterProfiler (39). Hallmark pathways were used. Genes upregulated in the QIIV + LVA group on day 1 were subjected to network analysis in Cytoscape (v3.10.4) (40).

#### Single cell RNA sequencing analysis

Targeted transcriptomics FASTQ files were processed via the standard Rhapsody analysis pipeline (BD Biosciences) on Seven Bridges (https://www.sevenbridges.com), according to the manufacturer’s recommendations. The filtered count matrix was then analyzed using the Seurat package in R (41). Low quality cells were removed. Doublet cells were removed from the analysis using the DoubletFinder package in R (42). Cell annotation was carried out using the reference expression dataset derived from Azimuth and further refined manually using the expression of known cell markers. Cells were clustered using the Louvain algorithm and visualized using Uniform Manifold Approximation and Projection (UMAP), and global differences between clusters were assessed using PCA. Differences between QIIV + LVA and QIIV in relative frequencies of different cell types at all timepoints were analyzed using Wilcoxon Rank test with correction for multiple testing (Benjamini-Hochberg procedure). DEA was performed using FindMarkers in the Seurat Package for the same conditions as bulk RNA sequencing (Days *and* Vaccine). Ranked DEGs per cell type were used for GSEA using the clusterProfiler package. Cell-Cell communication analyses were done using the R package CellChat (43).

### Statistical analysis

Categorical variables were summarized using counts and percentages, while continuous variables were presented as means, medians, standard deviations (SD), standard error of the mean (SEM), interquartile ranges (IQR), 95% confidence intervals (CI), minimum (min), and maximum (max). Differences in polypositive CD4+ T cell frequencies, serum cytokine concentrations and IgG isotype concentrations between cohorts at each visit were analyzed using Wilcoxon rank-sum test, with multiple testing correction via the Benjamini-Hochberg procedure. For correlation analyses, Spearman correlation coefficients were calculated. A p-value <.05 was considered statistically significant. All statistical analyses and data visualizations were performed using R and RStudio.

## Data availability

Raw data, including de-identified participant data, will be made available upon reasonable request to the corresponding author. Bulk RNA sequencing data is available at GSE336590 and single cell RNA sequencing data at GSE338106.

## Code availability

The full code to generate figures and perform analyses is available on Github.

## Acknowledgements

We thank all the participants in the study, and the clinical and laboratory staff of CEVAC. We would like to acknowledge the INDIGO Consortium (https://indigo-vaccines.eu/) in the generation of data used in this publication.

The INDIGO project, “Effective and Affordable flu Vaccine for the World”, is funded by the European Union Horizon 2020 programme under grant agreement No. H2020-SC1-2019-874653-10-INDIGO and the Department of Biotechnology, Ministry of Science and Technology, Government of India (BT/IN/EU-INF/17/GK/19-20). Views and opinions expressed are however those of the author(s) only and do not necessarily reflect those of the European Union or the Government of India. Neither the European Union nor the Government of India can be held responsible for them.

## Author contributions

Conceptualization: VDO, GLR, PPP, LH, ILR; Methodology: VDO, MV, SP, BJ, AA, SDG, SV, AW, GW, FDB, EDM; Formal analysis: VDO, MV, SP, SV, AW, EDM; Investigation: VDO, BJ, AA, SDG, AW, FDB, PPP, LH, ILR; Resources: VDO, GW, FDB, EVR, PPP, LH, ILR; Data curation: VDO, MV, SP, SV, AW, GW, LH; Writing (original draft): VDO; Writing (review & editing): all, Visualization: VDO, MV, SP; Supervision: GLR, EVR, PPP, LH, ILR; Project administration: VDO, MV, SP, GW, FDB; Funding acquisition: VDO, GW, FDB, EVR, PPP, LH, ILR.

## Figures

**Extended Data Figure 1.**
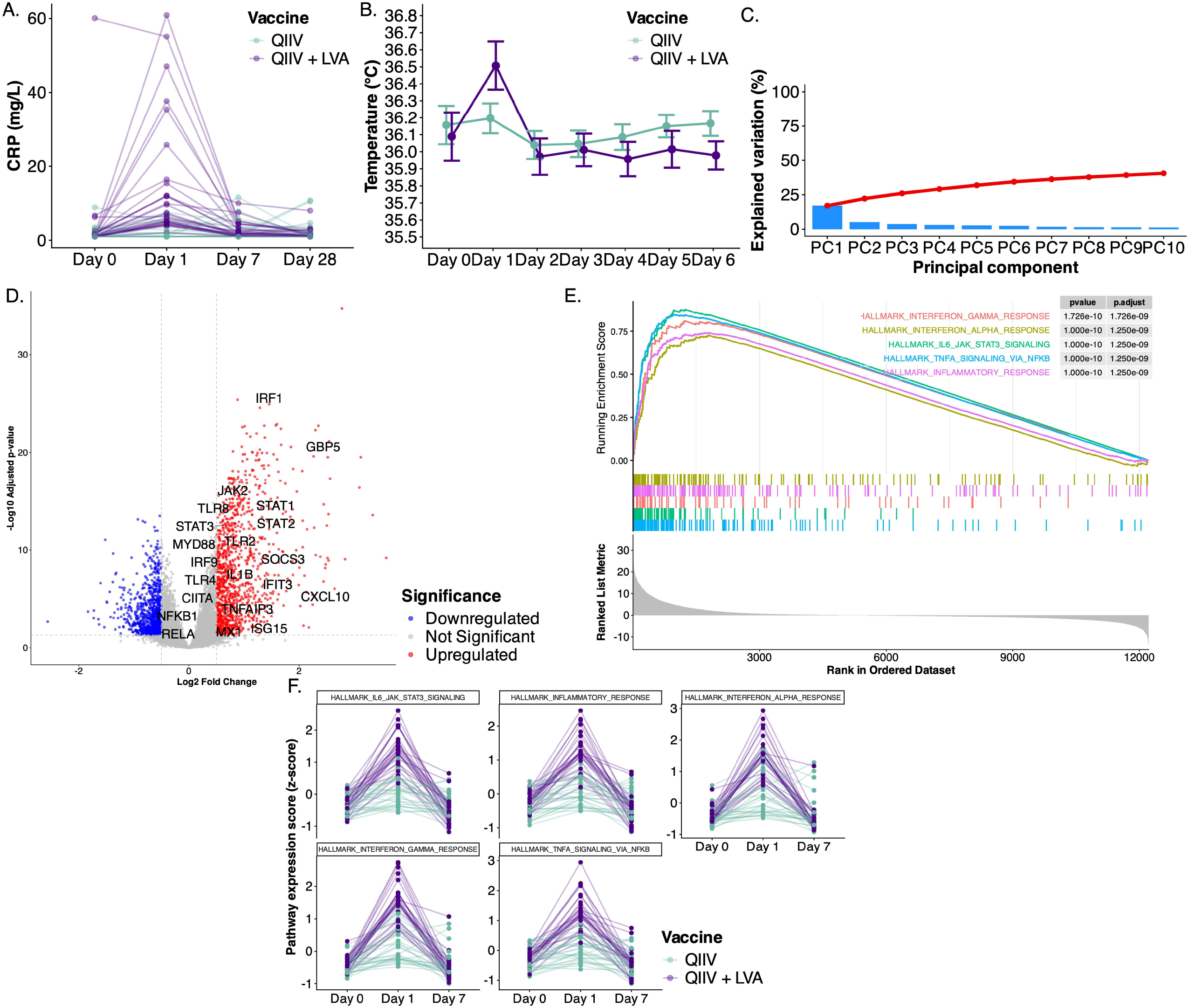
Clinical and transcriptomic correlates of early vaccine-induced inflammation. (A) Serum CRP concentrations per participant across time following vaccination. CRP increases transiently after immunization, peaking at early timepoints, with variability across individuals. (B) Mean + SEM of body temperature measurements over the first week post-vaccination. No sustained fever is observed, indicating that vaccination induces mild and self-limiting systemic responses. (C) Variance explained by principal components derived from bulk transcriptomic data. The first principal component captures the largest proportion of variance, consistent with a dominant vaccine-induced transcriptional program. (D) Volcano plot of differential gene expression at early timepoints post-vaccination relative to baseline. Significantly upregulated genes that are labelled include key innate immune and inflammatory regulators (e.g., CXCL10, IFIT3, ISG15, STAT1/2/3, IRF1/9, TLRs, MYD88, NFKB1), highlighting activation of IFN and antiviral pathways. (E) Gene set enrichment analysis (GSEA) of Hallmark pathways at day 1 post-vaccination between vaccine groups illustrating enrichment of interferon and inflammatory-related Hallmark pathways. Top enriched pathways include IFN*γ* and IFNα, responses with strong statistical significance after multiple-testing correction. (F) Temporal dynamics of pathway activity scores (z-scores) for selected Hallmark pathways in each participant stratified by vaccine group (QIIV vs. QIIV + LVA). Pathways associated with interferon signaling and inflammation are rapidly induced at day 1 and partially resolve by day 7, with generally stronger and more sustained activation in the QIIV + LVA group. LVA: Litevax Adjuvant; QIIV: quadrivalent inactivated seasonal influenza vaccine.

**Extended Data Figure 2.**
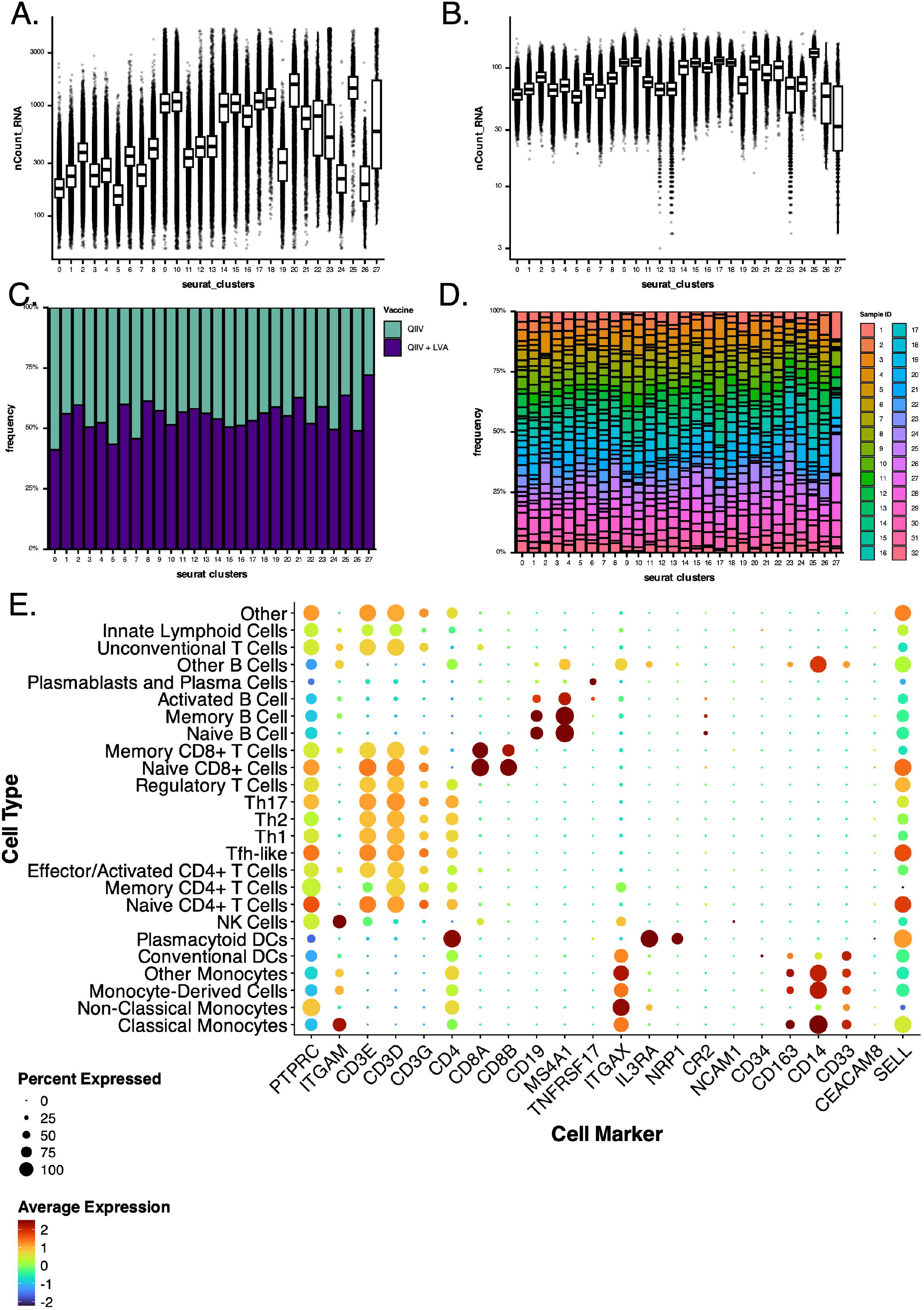
Single-cell RNA sequencing quality control, clustering and cell type annotation. (A) Distribution of total counts per cell (nCount_RNA) across clusters. (B) Distribution of detected genes per cell (nFeature_RNA) across identified single-cell clusters. (C) Proportion of cells derived from each vaccine group (QIIV vs. QIIV + LVA) across clusters. Stacked bar plots indicate comparable representation of both conditions across most clusters, supporting balanced sampling. (D) Composition of individual samples across single-cell clusters. Each bar represents a sample, colored by cluster identity, illustrating inter-individual variability and overall consistency in cluster distribution. (E) Dotplot of canonical marker gene expression used for cell type annotation. Dot size indicates the percentage of cells expressing each marker, and color intensity reflects average expression level. Distinct expression patterns define major immune populations, including monocyte subsets (CD14, FCGR3A), DCs (CLEC9A, LILRA4), T cell subsets (CD3D, CD4, CD8A, FOXP3), B cell populations (MS4A1, CD79A), and plasmablasts (MZB1). LVA: Litevax Adjuvant; QIIV: quadrivalent inactivated seasonal influenza vaccine.

**Extended Data Figure 3.**
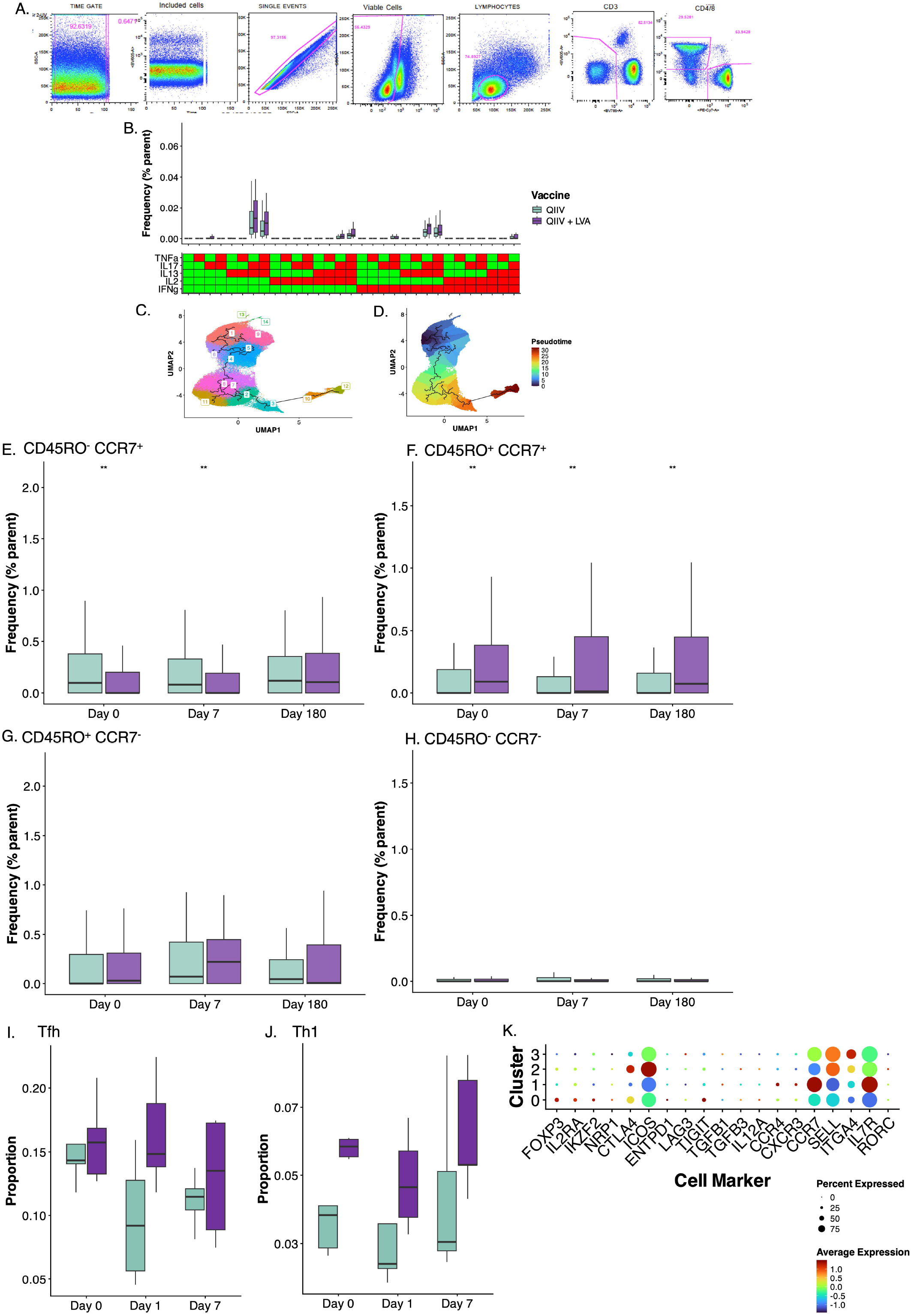
Functional and phenotypic characterization of CD4+ T cell responses. (A) Flow cytometry gating strategy used to identify antigen-specific CD4+ T cells and cytokine-producing subsets following stimulation. (B) Frequencies of cytokine-producing CD4+ T cells (including TNFα, IL-17, IL-13, and IFN*γ*) following vaccination. The QIIV + LVA group shows enhanced functional responses, particularly production of IFN*γ* and TNFα compared to QIIV alone. Green indicates positive for the marker, while red indicates negative for the marker. (C) UMAP projection of gated CD4+ T cells colored by cluster identified using unsupervised analysis. Data from intracellular cytokine staining (ICS) and flow cytometry. (D) UMAP projection of gated CD4+ T cells colored by pseudotime, highlighting transcriptionally distinct states across the differentiation landscape. Data from ICS and flow cytometry. (E-H) Frequencies of CD4+ T cell subsets defined by CD45RO and CCR7 marker positivity (flow cytometry) across time (days 0, 7, and 180). (E) Naïve (CD45RO^−^CCR7^+^), (F) central memory (CD45RO^+^CCR7^+^), (G) effector memory (CD45RO^+^CCR7^−^), and (H) terminal/effector-like (CD45RO^−^CCR7^−^). (I-J) Temporal changes in proportions of T follicular helper (Tfh) and Th1 cells in single-cell RNA sequencing data following vaccination, demonstrating induction of Th1 cells at day 7. (K) Dotplot showing expression of key markers for the annotation of Treg subsets, including regulatory (e.g., FOXP3, IL2RA, CTLA4, TIGIT), activation (e.g., ICOS), trafficking (e.g., CCR7, SELL), and lineage-defining transcriptional programs (e.g., RORC, CXCR3). Dot size represents the percentage of expressing cells and color indicates average expression. Significance levels are indicated as * (p < 0.05), ** (p < 0.01), *** (p < 0.001), **** (p < 0.0001), unless otherwise specified. LVA: Litevax Adjuvant; QIIV: quadrivalent inactivated seasonal influenza vaccine.

**Extended Data Figure 4.**
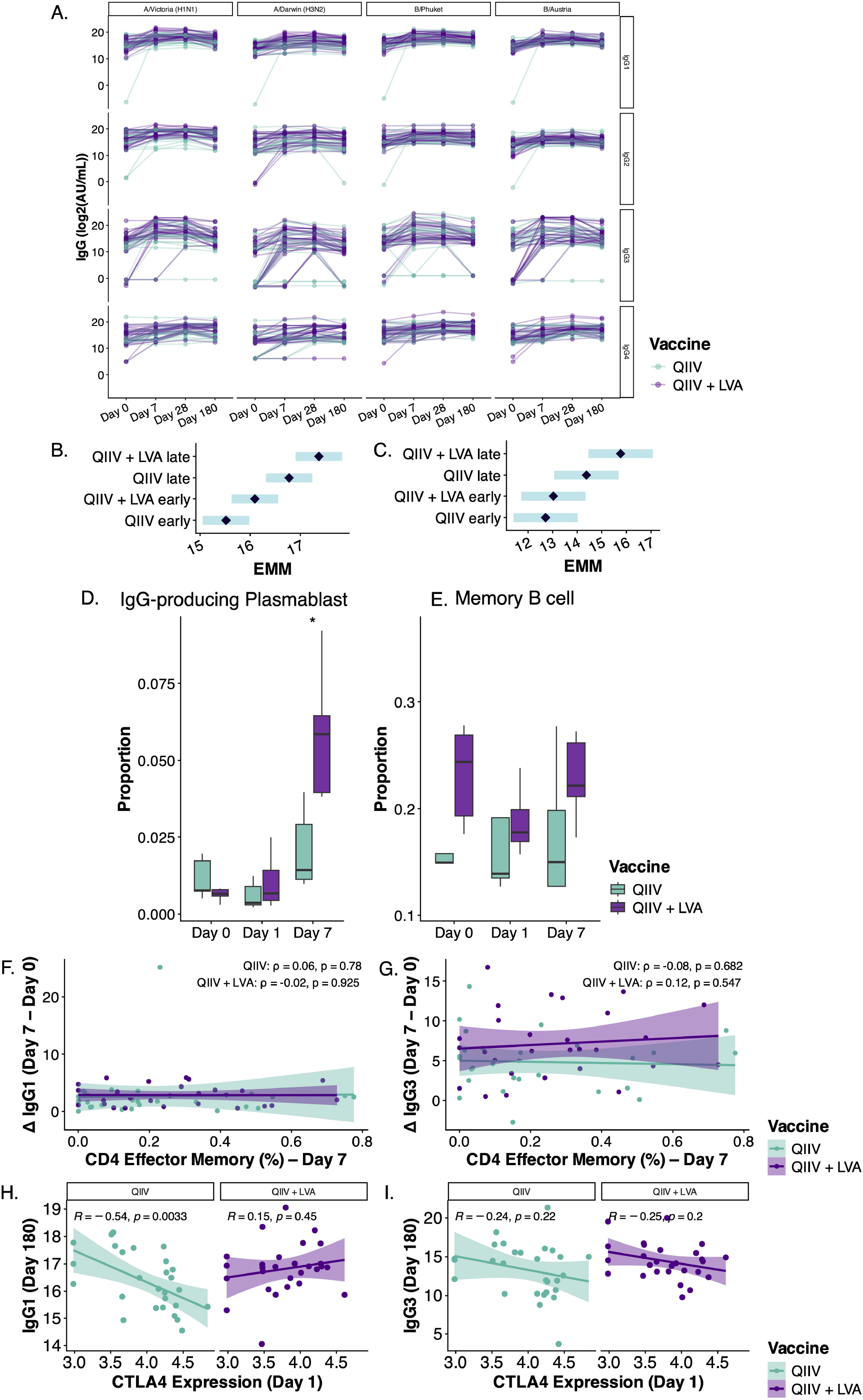
Relationship between B cell responses, antibody subclasses, and T cell help. (A) Longitudinal IgG subclass responses (IgG1, IgG2, IgG3, and IgG4) against individual influenza strains (A/Victoria H1N1, A/Darwin H3N2, B/Phuket, B/Austria) measured at days 0, 7, 28, and 180 per participant. Antibody kinetics reveal strain-specific and subclass-specific differences. (B-C) Estimated marginal means (EMMs) derived from linear mixed-effects models of IgG1 (B) or IgG3 (C) responses. EMMs summarize adjusted group means across early (day 0 and day 7) and late (day 28 and day 180) timepoints, accounting for repeated measures and inter-individual variability. Comparisons between QIIV and QIIV + LVA conditions show no differences in the magnitude and kinetics, but comparisons between timepoints show significant induction of IgG1 and IgG3 responses in late compared to early timepoints. (D) Proportions of IgG-producing plasmablasts over time (days 0, 1, and 7). Vaccination induces rapid expansion of plasmablast populations, with stronger early responses in the QIIV + LVA condition. (E) Proportions of memory B cells over time (days 0, 1, and 7). Vaccination does not induce significant changes in the proportions of memory B cells. (F-G) Correlation between CD4+ effector memory T cell frequencies (defined by flow cytometry: CD45RO^+^ CCR7^-^) at day 7 and (F) early IgG1 responses (ΔIgG1, day 7 minus day 0) or (G) IgG3 responses (ΔIgG3, day 7 minus day 0). No significant association is observed in either vaccine group. (H-I) Correlation between CTLA4 expression at day 1 (bulk RNA transcriptomics) and (H) long-term IgG1 responses (day 180) or (I) long-term IgG3 responses (day 180). A significant negative correlation is observed in the QIIV group but not in the QIIV + LVA group, suggesting differential regulation of antibody durability. Correlations are shown as Spearman’s R. Significance levels are indicated as * (p < 0.05), ** (p < 0.01), *** (p < 0.001), **** (p < 0.0001), unless otherwise specified. LVA: Litevax Adjuvant; QIIV: quadrivalent inactivated seasonal influenza vaccine.

**Extended Data Figure 5.**
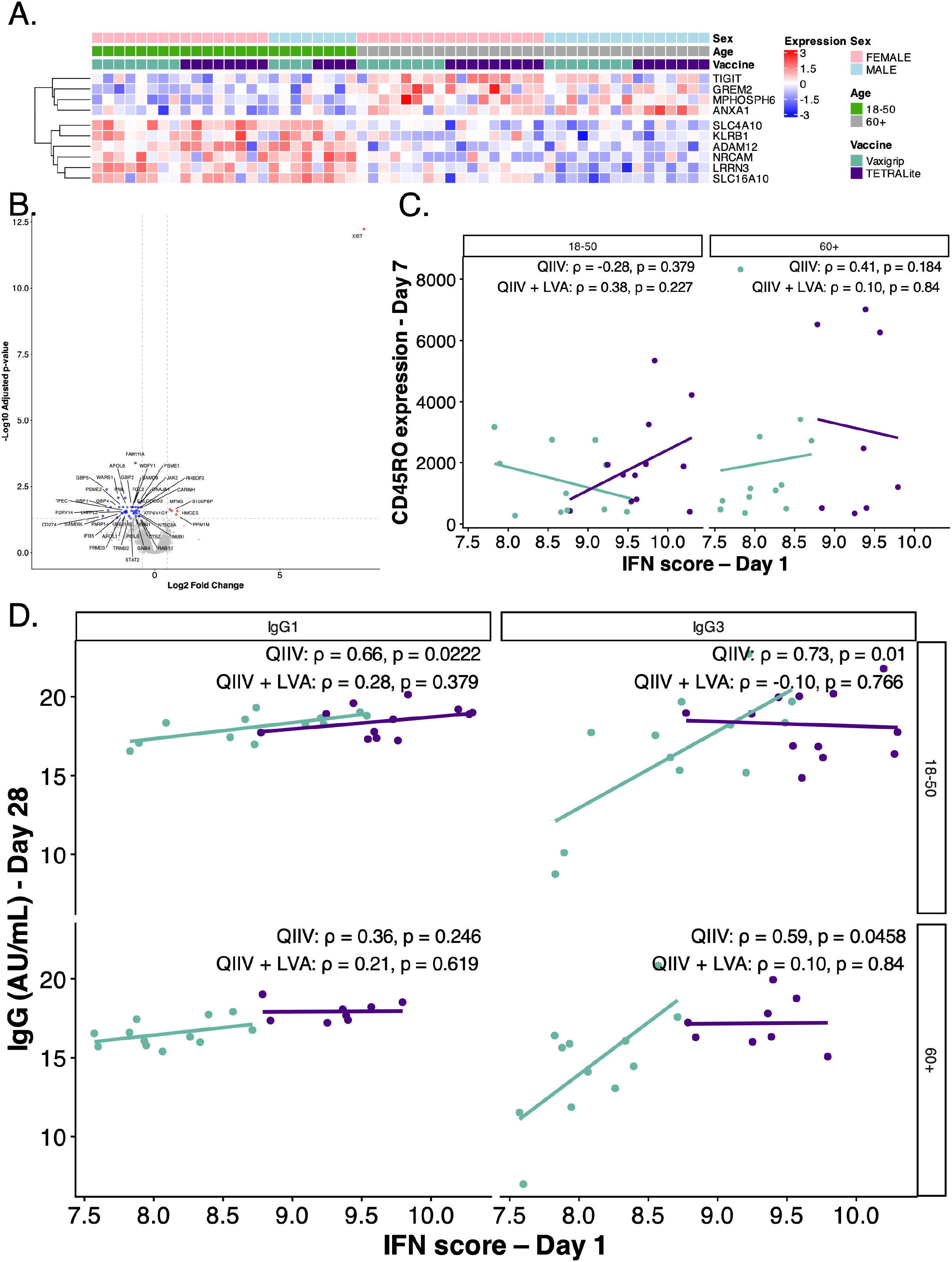
Age-associated transcriptional signatures. (A) Heatmaps showing expression of differentially expressed genes at baseline (day 0) between OA and YA. Genes include regulators of immune activation, signaling, and cellular metabolism (e.g., TIGIT, KLRB1, ANXA1, NRCAM), with distinct expression patterns highlighting age-dependent transcriptional programs. (B) Volcano plot of differential gene expression between OA and YA at day 1, highlighting significantly downregulated genes (blue) or upregulated genes (red) in OA. (C) Correlations between age-associated transcriptional score at day 1 and CD4+ T cell activation (CD45RO marker intensity at day 7). Associations differ between vaccine groups, suggesting altered coupling between early innate responses and T cell activation in the presence of LVA. (D) Correlations between age-associated transcriptional score at day 1 and antibody responses (IgG1 and IgG3) at day 28, stratified by age group (18–50 vs. ≥60 years) and vaccine condition (QIIV vs. QIIV + LVA). Strong positive associations are observed in the QIIV group, particularly in younger individuals, whereas these relationships are attenuated or absent in the QIIV + LVA group. Significance levels are indicated as * (p < 0.05), ** (p < 0.01), *** (p < 0.001), **** (p < 0.0001), unless otherwise specified. LVA: Litevax Adjuvant; OA: older adults; QIIV: quadrivalent inactivated seasonal influenza vaccine; YA: younger adults.

